# *TP53* loss initiates chromosomal instability in high-grade serous ovarian cancer

**DOI:** 10.1101/2021.03.12.435079

**Authors:** Daniel Bronder, Darawalee Wangsa, Dali Zong, Thomas J. Meyer, René Wardenaar, Paul Minshall, Anthony Tighe, Daniela Hirsch, Kerstin Heselmeyer-Haddad, Louisa Nelson, Diana Spierings, Joanne C. McGrail, Maggie Cam, André Nussenzweig, Floris Foijer, Thomas Ried, Stephen S. Taylor

## Abstract

High-grade serous ovarian cancer (HGSOC) originates in the fallopian tube epithelium and is characterized by ubiquitous *TP53* mutation and extensive chromosomal instability (CIN). While the direct causes of CIN are errors during DNA replication and/or chromosome segregation, mutations in genes encoding DNA replication and mitotic factors are rare in HGSOC. Thus, the drivers of CIN remain undefined. We therefore asked whether the oncogenic lesions that are frequently observed in HGSOC are capable of driving CIN via indirect mechanisms. To address this question, we genetically manipulated non-transformed *hTERT*-immortalized human fallopian tube epithelial cells to model homologous recombination deficiency (HRD) and oncogenic signalling in HGSOC. Using CRISPR/Cas9-mediated gene editing, we sequentially mutagenized the tumour suppressors *TP53* and *BRCA1*, followed by overexpression of the *MYC* oncogene. Single-cell shallow-depth whole-genome sequencing revealed that loss of p53 function was sufficient to lead to the emergence of heterogenous karyotypes harbouring whole chromosome and chromosome arm aneuploidies, a phenomenon exacerbated by subsequent loss of BRCA1 function. In addition, whole-genome doubling events were observed in independent p53/BRCA1-deficient subclones. Global transcriptomics showed that *TP53* mutation was also sufficient to deregulate gene expression modules involved in cell cycle commitment, DNA replication, G2/M checkpoint control and mitotic spindle function, suggesting that p53-deficiency induces cell cycle distortions that could precipitate CIN. Again, loss of BRCA1 function and MYC overexpression exacerbated these patterns of transcriptional deregulation. Thus, our observations support a model whereby the initial loss of the key tumour suppressor *TP53* is sufficient to deregulate gene expression networks governing multiple cell cycle controls, and that this in turn is sufficient to drive CIN in pre-malignant fallopian tube epithelial cells.

**SUMMARY STATEMENT:** High-grade serous ovarian cancer is defined by *TP53* mutation and chromosomal instability, the cause of which remains poorly understood. We developed a novel model system that implicates cell cycle deregulation upon p53-loss as cause of CIN.

## INTRODUCTION

High-grade serous ovarian cancer (HGSOC) is the most common histological sub-type of ovarian cancer, and the deadliest gynaecological malignancy (Bowtell et al., 2015). Survival statistics are dismal, with 5-year survival of ∼30%, and have remained largely unchanged over the past 30 years, illustrating the need for improved therapeutic interventions, which requires a better understanding of the underlying disease biology.

HGSOC is characterised by a relatively low mutational burden at the nucleotide level (Ciriello et al., 2013). *TP53* mutations are ubiquitous and are considered to be an early, truncal event in HGSOC tumorigenesis, which are present in precursor lesions (Ahmed et al., 2010; Labidi-Galy et al., 2017; Vang et al., 2016). However, with the exception of *BRCA1/2* mutations in ∼25% of cases, other common driver mutations are rare (Cancer Genome Atlas Research, 2011). By contrast, HGSOC genomes are characterized by extensive chromosomal copy number aberrations, a consequence of rampant chromosomal instability (CIN) (Cancer Genome Atlas Research, 2011; Nelson et al., 2020). Indeed, HGSOC ranks among the most chromosomally unstable tumour types (Ciriello et al., 2013; Shukla et al., 2020), a characteristic confirmed by recent live cell imaging of established cell lines and patient-derived *ex vivo* cultures, which revealed an unprecedented level of chromosome segregation errors (Nelson et al., 2020; Tamura et al., 2020).

To delineate the mechanisms responsible for CIN, HGSOC genomes have been extensively studied by whole genome sequencing, with one study defining two CIN classes, characterized either by homologous recombination deficiency (HRD) or foldback inversions (FBI) (Wang et al., 2017). While the former correlated with mutations in *BRCA1/2*, amplifications of *MECOM* and *MYC,* and loss of *RB1*, the latter correlated with *CCNE1* amplification and *PTEN* loss (Wang et al., 2017). A second study identified seven CIN signatures, including whole-genome duplication (WGD), suggesting a larger array of underlying driver mechanisms in addition to HRD and FBI (Macintyre et al., 2018).

This presents a paradox; while HGSOC appears to be driven by CIN, mutations in genes ensuring faithful cell division and DNA replication are extremely rare (Bastians, 2015). HRD, either as a consequence of *BRCA1/2* inactivation or mutation in other DNA damage repair genes is an obvious contributor to CIN, but by itself can only account for up to ∼50% of cases (Cancer Genome Atlas Research, 2011; Weaver et al., 2002; Xu et al., 1999). *TP53* has consistently been shown to correlate with aneuploidy (Ciriello et al., 2013; Davoli et al., 2017; Taylor et al., 2018; Zack et al., 2013), but its role as a driver of CIN remains controversial. Initial studies using the near-diploid colorectal cancer cell line HCT116, suggested that p53-loss is not sufficient to cause CIN (Bunz et al., 2002). More recently, however, suppressing p53 in *hTERT*-immortalized RPE-1 cells did generate abnormal karyotypes (Kok et al., 2020; Soto et al., 2017). Furthermore, p53 inactivation in transformed murine embryonic fibroblasts deregulated multiple cellular processes affecting DNA damage response, mitosis and ploidy control (Valente et al., 2020).

Here, we aimed to develop novel model systems of CIN in HGSOC, starting with *hTERT*-immortalized non-ciliated fallopian tube epithelial cells (Merritt et al., 2013). In the first instance, we set out to model the HRD CIN class, using CRISPR/Cas9-mediated gene editing to first mutate *TP53* then *BRCA1*, followed by overexpression of *MYC.* A panel of derivative subclones were subjected to functional assays, karyotyping and gene expression profiling to determine whether (a) CIN had been induced and (b) what the potential mechanisms might be.

## RESULTS

### FNE1 cells to model CIN in HGSOC

In addition to the truncal *TP53* mutation, *BRCA1/2* mutations and *MYC* overexpression tend to co-occur (Wang et al., 2017), suggesting that HRD and oncogene hyperactivation likely facilitate the development of CIN in HGSOC (Fig. 1A). To model these events, we set out to manipulate diploid, karyotypically stable cells, sequentially mutating *TP53* and *BRCA1* using CRISPR/Cas9-mediated gene editing, followed by ectopic overexpression of *MYC* (Fig. 1B). Since the fallopian tube epithelium is the likely origin for HGSOC we chose the human FNE1 cell line as a starting point (Ducie et al., 2017; Merritt et al., 2013). This line is derived from non-ciliated fallopian tube epithelial cells and immortalised by ectopic expression of the telomerase component *hTERT* (Merritt et al., 2013). Importantly, FNE1 cells are *TP53* proficient, evidenced by nuclear accumulation of p53 and p21 induction in response to the MDM2 inhibitor Nutlin-3 and to cisplatin (Fig. S1A,B and data not shown) (Vassilev et al., 2004). In addition, FNE1 cells are near-diploid and karyotypically stable, as confirmed by single-cell whole genome sequencing (scWGS) and spectral karyotyping (SKY). scWGS showed that the genome is largely disomic, except for monosomies at 9p, 15, and X (Fig. S1C). Consistently, SKY showed a clonal loss of chromosomes 15 and X and an unbalanced translocation between the short arm of chromosome 9 and chromosome 15 (Fig. S1D). An identical karyotype was also recently reported for FNE1 cells using multiplex fluorescence *in situ* hybridization (M-FISH) (Tamura et al., 2020). To enable CRISPR/Cas9-mediated gene editing in FNE1 cells, we transduced them with a lentivirus expressing a tetracycline-inducible Cas9 transgene. Increasing concentrations of tetracycline resulted in a dose-dependent induction of Cas9 (Fig. S1E). Importantly, in the absence of tetracycline, Cas9 was not detectable, thereby minimizing exposure of the genome to endonuclease activity during routine cell culture.

**Figure 1:**
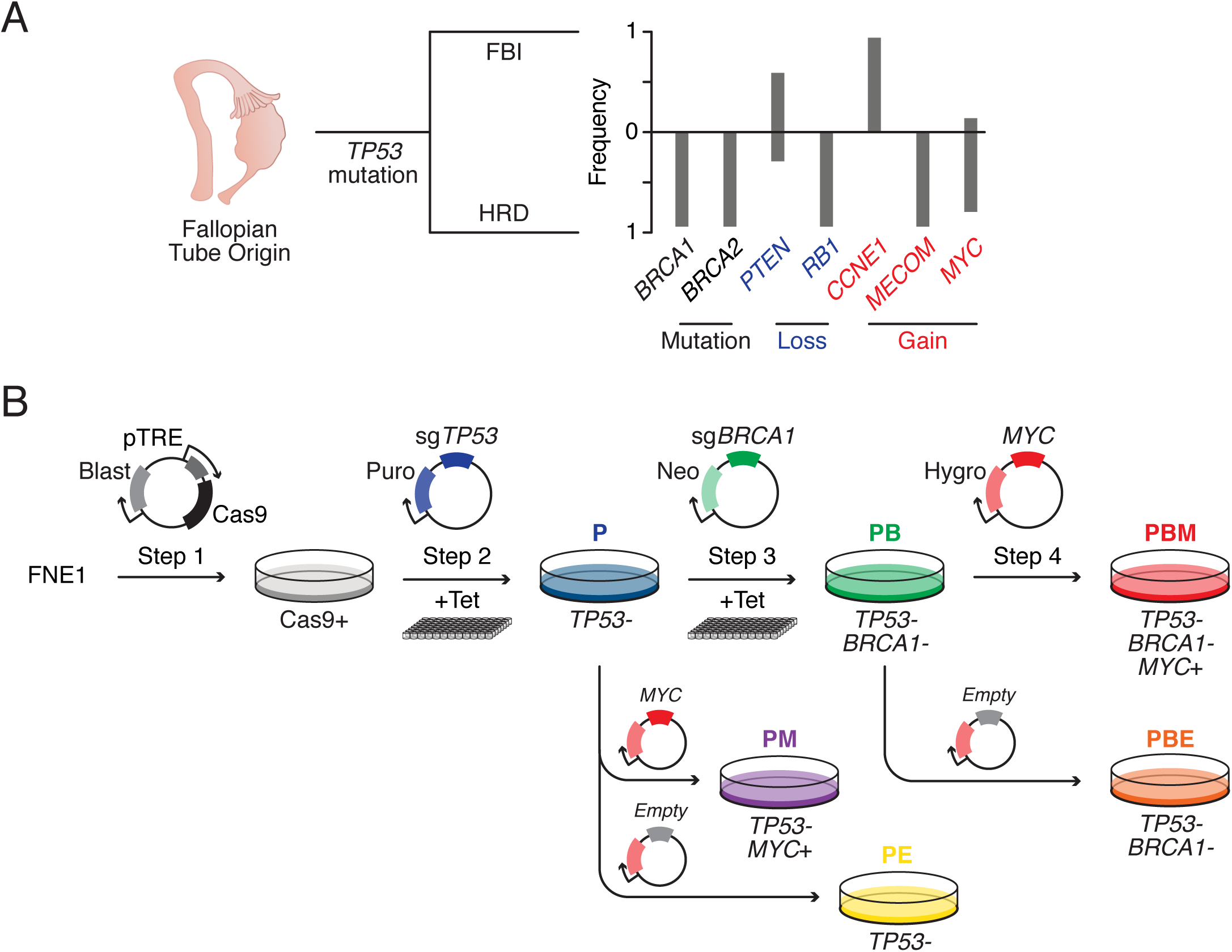
Intellectual Framework and Experimental Strategy. **A** Schematic of modelled high-grade serous ovarian cancer (HGSOC) development from the fallopian tube secretory epithelium including ubiquitous *TP53* mutation, grouping based on foldback inversions (FBI) or homologous recombination deficiency (HRD) and associated genomic changes in key tumour suppressors and oncogenes (Wang et al., 2017). **B** Experimental approach using *hTERT*-immortalized, fallopian tube-derived FNE1 cells to generate tet-inducible Cas9-expressing cells, which were then mutagenized to generate isogenic p53-deficient (P), p53/BRCA1-deficient (PB) and MYC-overexpressing double- (PM) and triple-(PBM) mutant subclones. MYC-overexpressing cells are co-isogenic, polyclonal populations of the parental subclones. Single- (PE) and double-mutant (PBE) control cells were also generated via transduction with an ‘*empty’* virus vector. See also Figure S2A.

### CRISPR/Cas9-mediated mutation of *TP53* and *BRCA1*

To mutate *TP53*, we introduced an sgRNA targeting exon 2, induced Cas9 then isolated subclones by limiting dilution, either with or without Nutlin-3 selection (Fig. 1B). Characterisation of three independent subclones, designated P1–3 (Fig. S2A, Table 1), showed an absence of p53 protein (Fig. 2A), and interrogation of RNAseq data showed that all three clones harboured frameshift mutations leading to premature termination codons (Table 1; Fig. S2B). Importantly, Nutlin-3 did not exert an anti-proliferative effect in the *TP53* mutants (Fig. 2B), indicating that the subclones are indeed functionally p53-deficient.

**Table 1.**
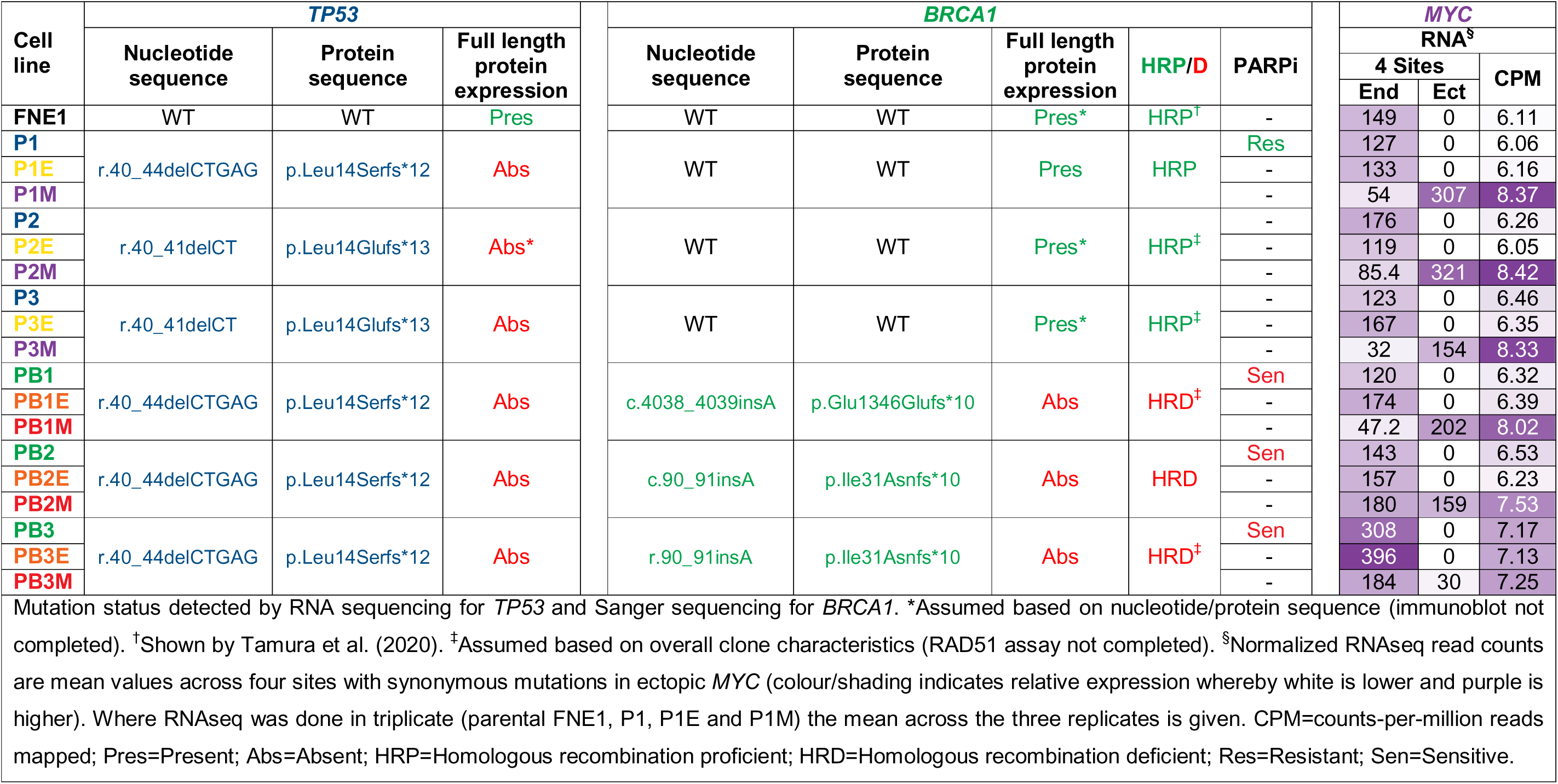
Summary of mutant cell lines generated in this study including mutation status and *MYC* RNA levels.

**Figure 2:**
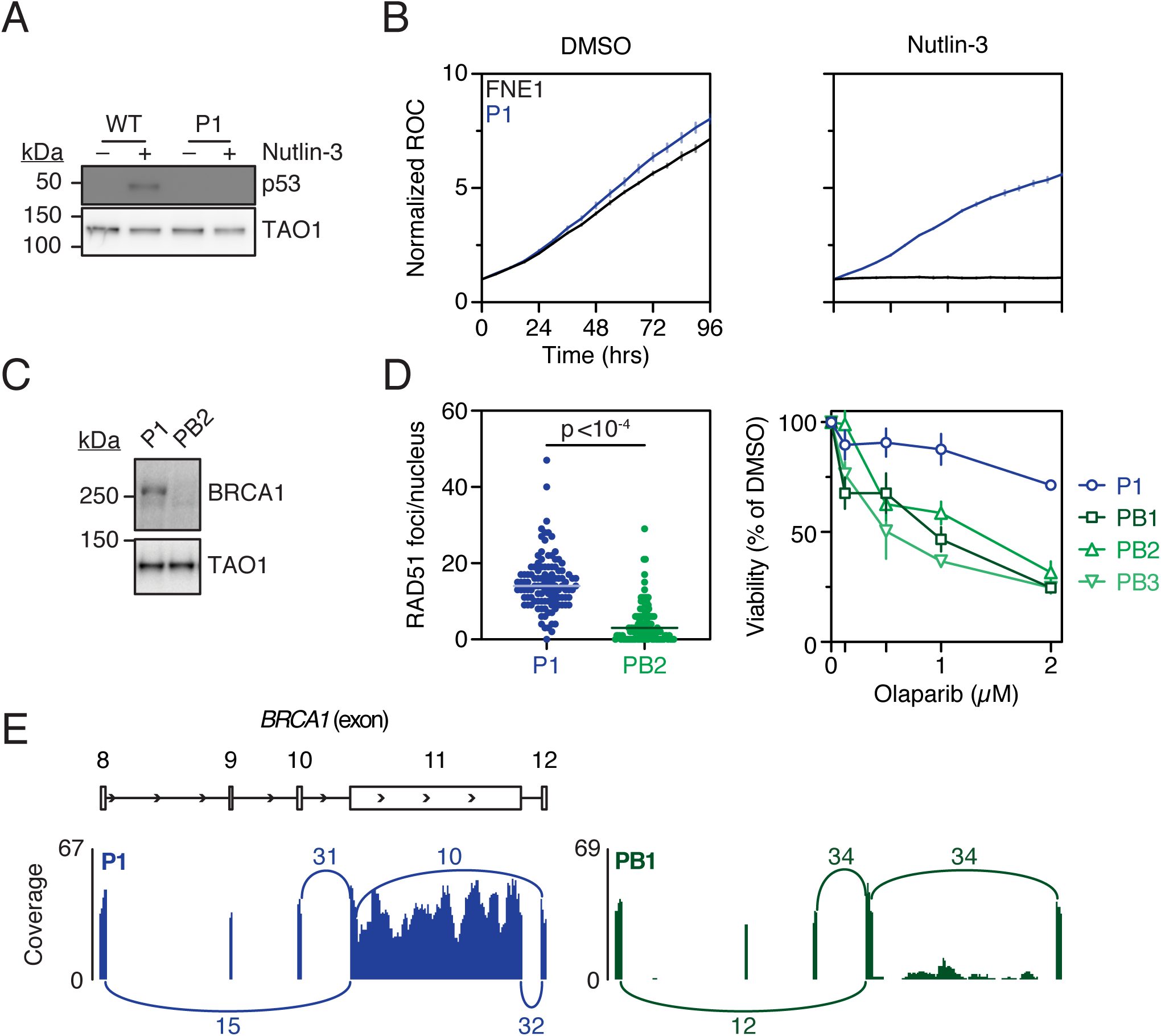
Generation and Functional Validation of *TP53* and *TP53*/*BRCA1*-mutant Subclones. **A** Representative immunoblot of p53 expression in CRISPR/Cas9-derived *TP53*-mutant (P1) cells and parental FNE1 cells treated with either DMSO (vehicle) or Nutlin-3. TAO1 serves as loading control. **B** Nuclear proliferation curves of parental FNE1 and P1 cells expressing an mCherry-tagged histone in the presence of DMSO or Nutlin-3. Normalised red object count (ROC) was calculated as fold change from T_0_. Results from three technical replicates are shown as mean with error bars indicating standard deviation. **C** Representative immunoblot of full-length BRCA1 expression in CRISPR/Cas9-derived *TP53/BRCA1* double-mutant (PB2) cells. Here, P1 reflects a BRCA1-proficient (p53-deficient) subclone recovered after Cas9 induction. TAO1 serves as loading control. **D** Left panel, Quantitation of RAD51 foci formation in EdU-positive *TP53*-mutant (P1; 111 nuclei) and *TP53*/*BRCA1* double-mutant (PB2; 114 nuclei) cells following 5 Gy ionizing radiation. Results from single experiment are shown. Statistical analysis was performed using a student’s t-test. Right panel, CellTiter-Blue® viability assay of P1 and PB1–3 cells treated with indicated concentrations of the PARPi olaparib over the course of one week. Viability was normalized to DMSO (vehicle)-treated cells. Results from three technical replicates, error bars represent standard deviation. **E** Representative Sashimi plot depicting alternative splicing of *BRCA1* exon 11 observed in P1 and PB1 subclones. Numbers indicate raw junction reads attesting to the splice events indicated by the arcs. The minimum of splice junction reads was three. Note that junction reads mapping 3’ terminally of exon 11 and 5’ terminally of exon 12 in PB1 are not detected in PB1. See also Figures S1, S2 and Table 1.

To then mutate *BRCA1*, clone P1 was transduced with sgRNAs targeting exons 2, 3 and 11 (Fig. S2A), Cas9 induced and subclones isolated by limiting dilution (Fig. 1B). Again, we characterised three independent subclones, designated PB1–3 (Table 1). Consistent with *BRCA1* mutation, immunoblotting failed to detect full length protein (Fig. 2C), induction of RAD51 foci in response to ionizing radiation was suppressed, and sensitivity to the PARP inhibitor olaparib was increased (Fig. 2D). To define the nature of the *BRCA1* mutations, we interrogated RNAseq data and mutations identified were then confirmed by Sanger sequencing of cloned genomic DNA (Table 1; data not shown). This revealed that PB2 and PB3 harboured mutations in exon 3, while PB1 harboured a mutation in exon 11. Interestingly, we observed alternative splicing of exon 11 in PB1 (Fig. 2E), an event that may lead to the production of a truncated BRCA1 protein that retains partial function (Wang et al., 2016). Thus, although all three PB subclones harbour *BRCA1* mutations, PB1 may have the capacity to retain partial homologous recombination (HR) proficiency. Altogether, these observations confirm the successful generation of FNE1 subclones harbouring mutations in both *TP53* and *BRCA1*.

### Ectopic overexpression of *MYC*

Following mutation of *TP53* and *BRCA1*, we set out to overexpress *MYC*, an oncogene frequently amplified in HGSOC. Indeed, across 18 tumour types, HGSOC displays the highest frequency of *MYC* amplification (Zeng et al., 2018). The three *TP53* mutant clones, P1–3, and the three P1-derived *TP53*/*BRCA1* double mutant clones, PB1–3, were all transduced with a lentivirus harbouring a *MYC* cDNA downstream of a constitutive CMV promoter, generating six polyclonal derivatives, designated P1–3M and PB1–3M (Fig. 1B, Fig. S2A). In parallel, we transduced an ‘*empty’* vector control virus, generating a further six polyclonal derivatives, designated P1–3E and PB1–3E (Fig. S2A). Note that the *MYC* cDNA harboured four synonymous mutations (Littler et al., 2019), allowing us to differentiate ectopic and endogenous *MYC* transcripts. In turn, RNA sequencing revealed that ectopic *MYC* was indeed overexpressed relative to endogenous *MYC* in P1–3M and PB1M (Fig. 3A). In PB2M and PB3M, however, the situation was reversed, possibly indicating endogenous *MYC* was already overexpressed in these two lineages. Indeed, *MYC* was highly expressed in PB3 and PB3E, consistent with spontaneous upregulation prior to our efforts to experimentally overexpress *MYC* (Table 1). However, for the PB2 lineage, *MYC* levels were only elevated in PB2M as expected following ectopic *MYC* overexpression, and not in PB2 or PB2E.

**Figure 3:**
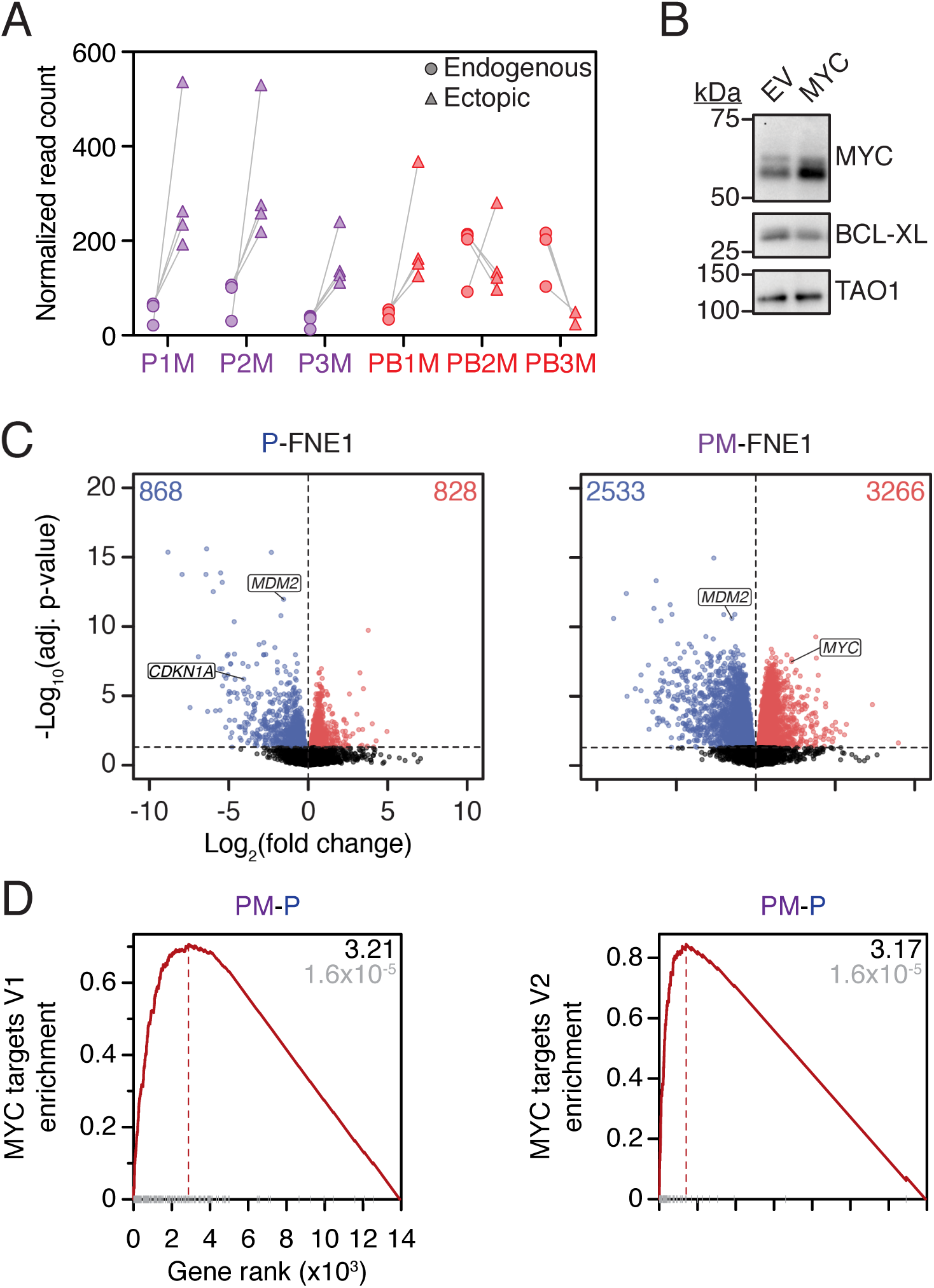
Generation and Functional Validation of *MYC*-overexpressing *TP53*-mutant and *TP53/BRCA1*-mutant Subclones. **A** Normalized read count of endogenous (circle) and ectopic (triangle) *MYC* RNA was determined by interrogating RNA sequencing data at the nucleotide level. Read counts at four sites of synonymous mutations in ectopic *MYC* were enumerated, with each mutation site reflected by one of the four circles/triangles per cell line in the graph. Reads were normalized to uniquely mapped reads. P1M was sequenced in triplicate thus the average of the three replicates is plotted for each locus. Note, endogenous *MYC* levels may be elevated in PB2M and PB3M relative to other samples (see results text). **B** Representative immunoblot of P1 cells transduced with empty vector (EV) or MYC-overexpressing (MYC) lentiviruses showing MYC and BCL-XL expression. TAO1 serves as loading control. **C** Volcano plots showing differentially expressed genes in P (pooled P1–3 and P1–3E) and PM (pooled P1–3M) samples, compared with parental FNE1 samples. Each point reflects a single gene where blue indicates differential down-regulation and red indicates differential up-regulation. Black means that the significance threshold of adj. p-value ≤ was not reached. The canonical p53 target genes *CDKN1A* and *MDM2* as well as *MYC* are indicated. The number of differentially down- and up-regulated genes is shown in blue and red font, respectively. **D** Enrichment of Hallmark MYC Targets V1 and V2 comparing PM (pooled P1–3M) with P (pooled P1–3 and P1–3E). Black font indicates normalized enrichment score, and grey font indicates adj. p-value. The adj. p-value for differentially expressed genes in C–D was determined using the Benjamini-Hochberg algorithm. Results are from a single experiment with pooled clones as described (with the exception of parental FNE1, P1, P1E and P1M, for which 3 technical replicates are included). P=*TP53*-mutant; B=*BRCA1-*mutant; E=Empty vector lentivirus; M=*MYC*-overexpressing lentivirus. See also Figure S2 and Table 1.

Importantly, overexpression of *MYC* modulated MYC-dependent processes, evidenced by immunoblotting of P1M cells, which revealed downregulation of the pro-survival factor BCL-XL (Fig. 3B). Consistent with MYC’s role as a transcriptional amplifier (Lin et al., 2012; Nie et al., 2020; Nie et al., 2012), analysis of differentially expressed genes in pooled P and PM cells revealed more significantly upregulated and downregulated genes upon overexpression of *MYC* (Fig. 3C). Moreover, gene set enrichment analysis (GSEA) showed that *MYC* hallmark target gene sets V1 and V2 are positively enriched in pooled PM cells versus controls (Fig. 3D). Interestingly, the V1 and V2 sets are also positively enriched versus parental FNE1 cells in both the PB2 and PB3 lineages, with and without introduction of ectopic MYC (see below; Fig.S5). Therefore, whilst PB3 lineage cells have likely enriched V1 and V2 sets via direct overexpression of endogenous *MYC*, PB2 lineage cells may have also spontaneously upregulated MYC target gene expression via an alternative mechanism, for example by alteration of downstream MYC signalling as has been observed previously in HGSOC samples (Jimenez-Sanchez et al., 2020). Thus, these observations confirm successful upregulation of *MYC* activity in FNE1 subclones harbouring mutations in *TP53* and *BRCA1*.

### Ploidy analysis reveals independent WGD events

Having established a panel of 18 FNE1 subclones harbouring genetic features found in HGSOC cells (Fig. S2A, Table 1), we set out to determine whether any of those displayed evidence of CIN. First, we analysed the P1 lineage by flow cytometry to explore changes in ploidy. The *TP53* mutant P1E, the *TP53*/*BRCA1* double mutant PB1E, plus their MYC-overexpressing counterparts, P1M and PB1M displayed typical 2c and 4c peaks, indicating no overt deviation from normal ploidy (Fig. S3). By contrast, the *TP53*/*BRCA1* double mutants, PB2E and PB3E, and their MYC-overexpressing counterparts, PB2M and PB3M, displayed evidence of 8c peaks, indicating a cycling tetraploid cell population. In PB2E and PB2M, the 8c peak was small and accompanied by 2c and 4c peaks, suggesting that only a sub-fraction of the population was tetraploid. While in PB3E and PB3M, the 4c and 8c peaks were more apparent than in PB2E/M and an obvious 2c peak was absent, suggesting that the entire population was tetraploid, i.e., had undergone WGD.

Because P1E and P1M appeared overtly normal, mutation of *TP53* alone or in combination with overexpression of *MYC* is not sufficient to induce tetraploidization. Moreover, the presence of tetraploidy in PB2E and PB3E also suggests that it arose prior to *MYC* overexpression. Rather, the flow cytometry suggests that the *BRCA1* mutation was possibly driving the tetraploidy. And yet, PB1E and PB1M, which also harbour *BRCA1* mutations, do not show evidence of tetraploidy. Note, however, that, as described above, we observed alternative splicing of exon 11 in PB1, raising the possibility that the BRCA1-deficiency in this line may not be as penetrant as in PB2 and PB3 lineages. Nevertheless, the presence of tetraploid cells in the PB2 and PB3 lineages suggests independent WGD events in*TP53*/*BRCA1* double mutant FNE1 cells.

### miFISH confirms WGD and reveals CIN

To obtain a more detailed picture of the ploidy changes observed by flow cytometry, we analysed 20 genetic loci in 100 FNE1, PB2M and PB3M cells using multiplex, interphase fluorescence *in situ* hybridization (miFISH) (Heselmeyer-Haddad et al., 2012). In parental FNE1 cells, 19 of the 20 loci analysed were predominantly present in two copies (Fig. 4A,C), consistent with a diploid and stable genome, and in line with the scWGS and SKY analysis (Fig. S1). In seven cells, we observed minor abnormalities, with one or two loci deviating from the mode; this, however, is within the margin of error of miFISH performed on cultured cells (Wangsa et al., 2018). By contrast, in every cell analysed only a single *CDKN2A* signal was detected, indicating a clonal loss of a region on chromosome 9, consistent with the karyotyping described above (Fig. S1). Note that the *CDKN2A* locus, which encodes the tumour suppressors p16 and p14ARF, is frequently altered in established cell lines, and may contribute to their unlimited proliferative potential *in vitro* (Huschtscha and Reddel, 1999).

**Figure 4:**
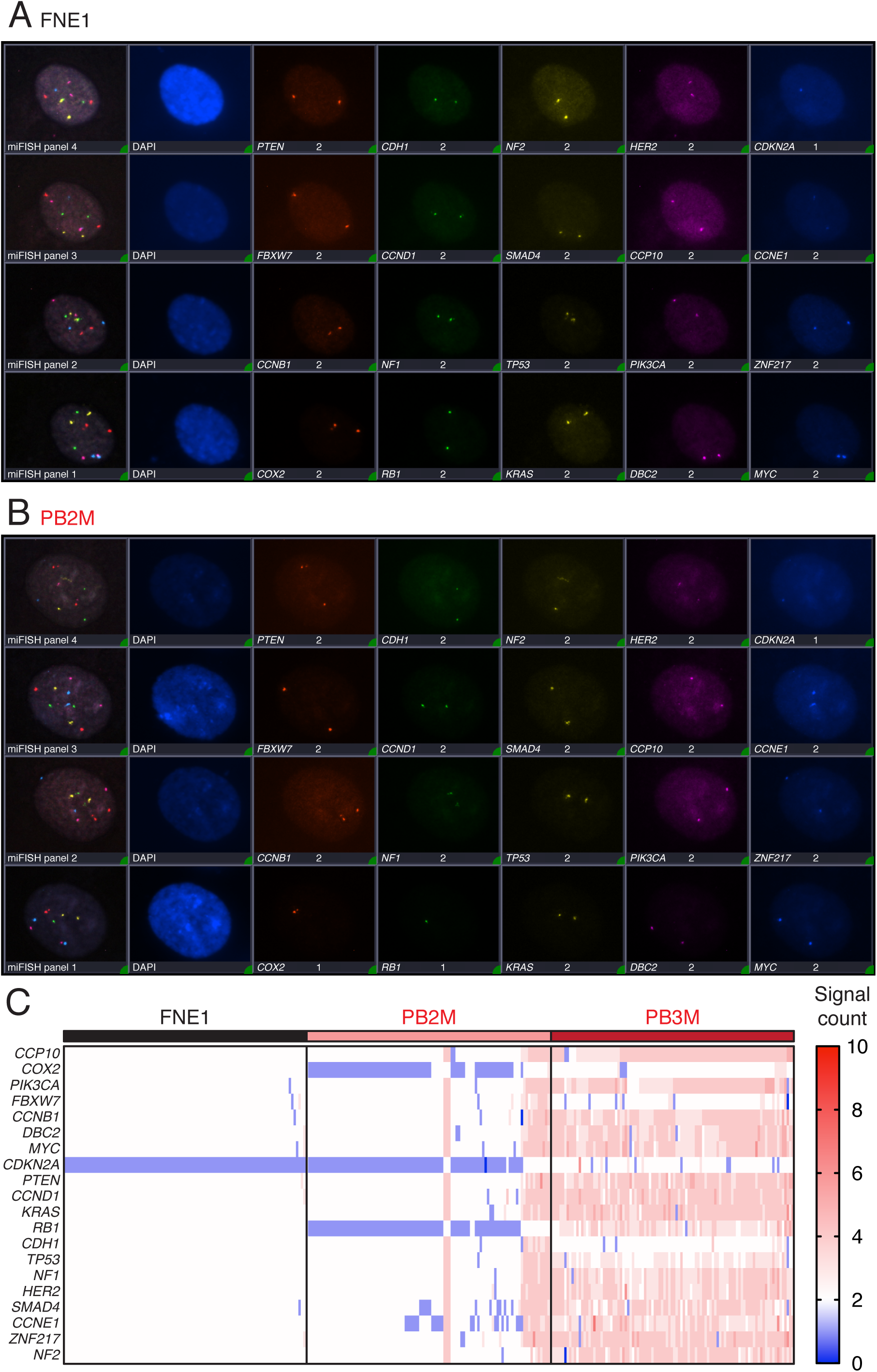
miFISH Implicates On-Going Chromosomal Instability, Aneuploidy and Whole Genome Doubling in Two Triple Mutant Subclones. **A–B** Representative composite multiplex, interphase fluorescence *in situ* hybridization (miFISH) images of all 20 probes hybridized in succession on parental FNE1 and PB2M cells, respectively. Note the reduced signal count of *COX2* and *RB1* in PB2M versus parental FNE1. **C** Copy number aberrations of centromere 10 (CCP10) and 19 indicated gene loci in parental FNE1 and the two aneuploid triple-mutant subclones assessed by miFISH. Blue and red indicate copy number loss and gain, respectively, relative to the diploid, parental FNE1. Columns indicate single cells (n=100, each for parental FNE1, PB1M and PB3M). P=*TP53*-mutant; B=*BRCA1*-mutant; M=*MYC*-overexpressing lentivirus. See also Figure S3.

In contrast to parental FNE1 cells, PB2M and PB3M displayed numerous deviations. As the ploidy measurements by flow cytometry suggested, PB2M harboured both 2c and 4c cells. The 2c subpopulation had the same clonal loss of *CDKN2A*, with additional clonal losses of *COX2* and *RB1* (Fig. 4B,C). These three clonal losses were also present in the 4c subpopulation, with only two foci of each detected. As expected, PB3M was confirmed by miFISH to be entirely composed of 4c cells (Fig 4.C). Like 4c PB2M cells, PB3M cells also had only two signals for some loci, i.e., *COX2*, *FBXW7*, *CDKN2A* and *CDH1*. These losses suggest that either a 4c population of PB3M cells has lost 2 copies of *COX2*, *FBXW7* and *CDH1*, but not *CDKN2A* (since its baseline is monosomic) or an elusive 2c PB3M population has undergone WGD; we favour the latter explanation. Interestingly, PB3M cells show a pattern of dosage decrease of chromosome 17. In most cells three copies of *TP53* were detected and four copies of *NF1* and *HER2*. In a subset where only two *TP53* signals were observed, three copies of *NF1* and *HER2* are seen. Overall, a more diverse pattern of gains and losses were detected in PB2/3M than in FNE1 cells. Thus, these observations confirm independent WGD events in lineages PB2 and PB3. Moreover, the sub-clonal gains and losses in both diploid and tetraploid backgrounds indicate the acquisition of CIN.

### scWGS reveals CIN in both diploid and tetraploid backgrounds

The sub-clonal gains and losses revealed by miFISH indicate CIN in the PB2M and PB3M lines. To explore this in more detail across a wider range of lines, and in particular in an unbiased, genome-wide manner, we performed scWGS-based karyotyping. In addition to parental FNE1 cells, we analysed the *TP53* mutant P1, the two *BRCA1*-deficient derivatives, PB2 and PB3, their *MYC*-expressing subclones, PB2M and PB3M, and the corresponding empty vector controls, PM2E and PB3E (Fig. S2A). Unsupervised hierarchical clustering identified four karyotype clusters (Fig. 5A). Cluster 1, which exhibited the monosomies at 9p, 15, and X described above (Fig. S1), consisted of parental FNE1 cells and the *TP53* mutant P1. Closer inspection revealed a number of partial or whole chromosome aneuploidies in P1 cells; whereas only two of 35 parental FNE1 cells (5.7%) displayed deviations, 10 of 18 P1 cells did so (55.6%), indicating that low level CIN is already present in *TP53*-deficient P1 cells.

**Figure 5:**
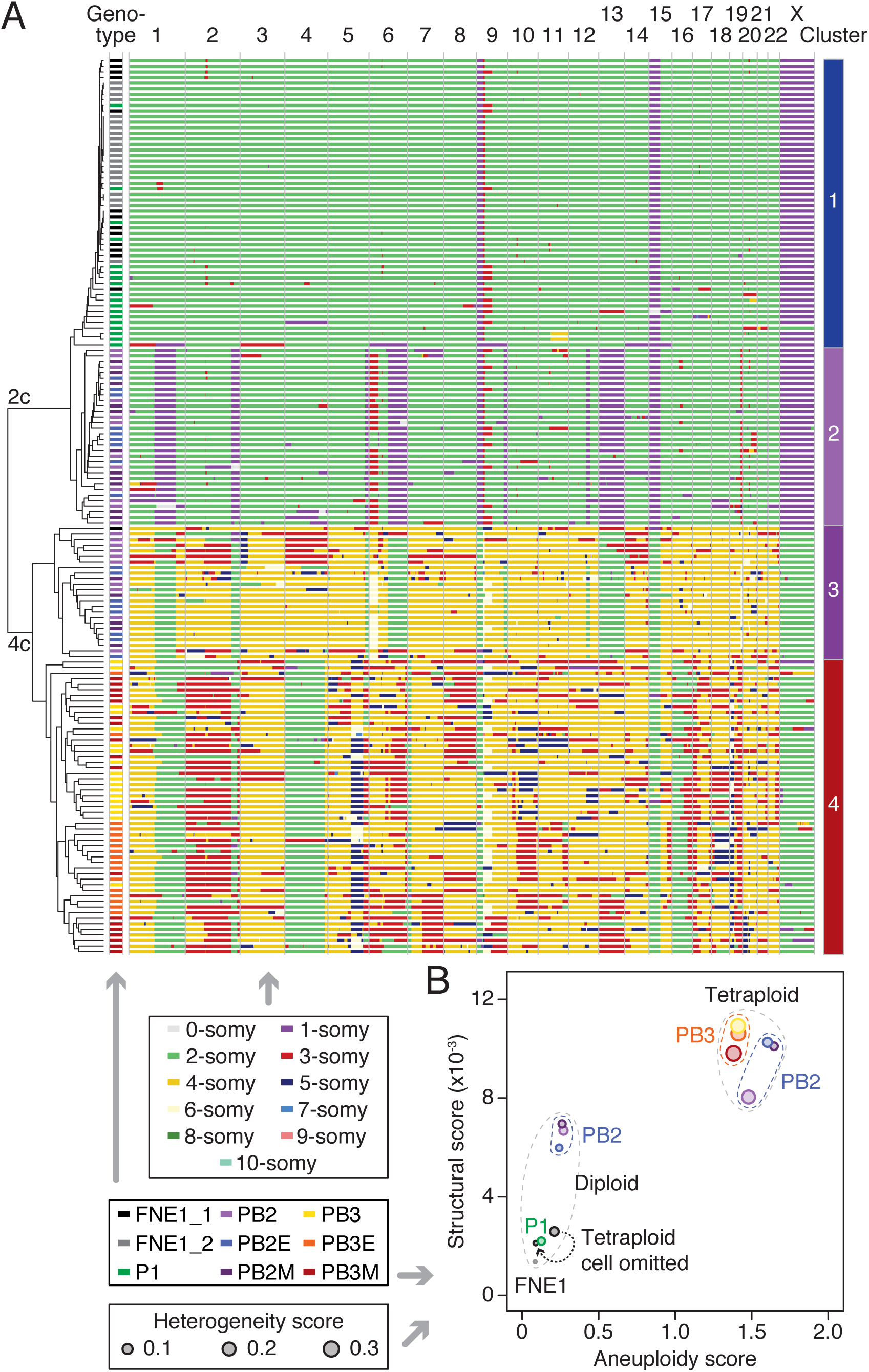
Single-cell Shallow-depth Whole-genome Sequencing Finds Ongoing CIN and Whole-Genome Doubling in Mutant Subclones. **A** Single cells from indicated genetic backgrounds were subjected to scWGS and subsequent unsupervised hierarchical clustering analysis, which first clusters cells by ploidy and then in a genotype-dependent manner. Autosomes from 1–22 and the X chromosome are displayed as columns. Each row represents a single cell of indicated genetic background (middle box). The colour in each row at a defined genomic location indicates the copy number (top box). Note FNE1_2 is a reproduction of data from Fig. S1C. **B** Aneuploidy, structural and heterogeneity scores were calculated from scWGS data in A. The structural score is defined as the number of copy number state transitions (within a single chromosome) per Mb, normalized to the number of cells analysed. Generation of the heterogeneity and aneuploidy scores are described previously (Bakker et al., 2016). Based on structural and aneuploidy scores samples separate into a diploid and tetraploid cluster. Note, one of the parental FNE1 samples contained a tetraploid cell (FNE1_1), which resulted in an increase in all three scores, which was reduced if the scores were re-calculated omitting that cell (dotted arrow).

Cluster 2 is characterised by near-diploid genomes with clonal segmental copy number losses on chromosomes 1, 2, 6, 12 and 13, a segmental gain on chromosome 6, and a variety of sub-clonal gains and losses. By contrast, cluster 3 was dominated by tetrasomies but with segmental disomies on chromosomes 1, 2, 6, 12 and 13, and various sub-clonal deviations. All the cells in clusters 2 and 3 were from the *TP53*/*BRCA2* double mutant lineage PB2, including PB2 itself, PB2M and PB2E, and thus reflect the diploid and tetraploid populations identified by miFISH analysis of PB2M. These data also corroborate the *COX2* (1q) and *RB1* (13q) losses seen in PB2M by miFISH, since the corresponding chromosome arms are monosomic in the diploid population. Importantly, because the monosomies in the diploid subpopulation are reflected as disomies in the tetraploid sub-population, these losses likely occurred prior to the WGD event. The increasing frequency of sub-clonal deviations in the diploid and tetraploid PB2-lineage populations (68.8% and 78.3% displaying deviations, respectively) compared with P1 indicates exacerbation of the low-level CIN induced by *TP53* loss.

Cluster 4, which is also dominated by tetrasomies, is made up exclusively of cells from the PB3 lineage, including PB3 itself, PB3M and PB3E, reflecting the tetraploid population identified by miFISH analysis of PB3M. Chromosomes 1q, 4 and 16 are disomic, suggesting clonal loss prior to WGD, while many other chromosomes display sub-clonal whole or segmental gains and losses, indicating pervasive CIN. Indeed, chromosome 5q displays features of rearrangement, loss and amplification. One particular segment is detectable as tetra-, penta- and hexasomy while the most telomeric region is present as di-, tri- and tetrasomy. A similar observation is made on chromosome 19 where 19p is predominantly detected in five or six copies and 19q is detected most frequently in three copies. Therefore, heterogeneity in the PB3 lineage also indicates that loss of BRCA1 function exacerbated low-level CIN induced by *TP53* loss.

### CIN is initiated by *TP53* loss and exacerbated by *BRCA1* mutation

Taking together, the ploidy analysis, the miFISH and the scWGS data, our observations support a model whereby, in the PB2 and PB3 lineages, *TP53* mutation initiated low-level CIN on an otherwise diploid background, which was then exacerbated by *BRCA1* mutation, followed by genome doubling events leading to tetraploidy and more pervasive CIN. While both diploid and tetraploid sub-clones are present in the PB2 lineage, the PB3 lineage is exclusively tetraploid, possibly reflecting an early WGD event during the genesis of this line. Importantly, the extensive CIN generated in our model system is reflective of M-FISH and scWGS from patient-derived *ex vivo* HGSOC cultures, which display profound inter-cellular heterogeneity with karyotypes characterized by whole-chromosome aneuploidies, rearranged chromosomes, monosomies and tetrasomies (Nelson et al., 2020).

While we did not observe CIN in the PB1 lineage, we did not perform scWGS so may have missed low-level deviations due to *TP53* loss. Also, due to alternative splicing of exon 11, this lineage may retain partial *BRCA1* function, explaining why more pervasive CIN did not manifest. Interestingly, overexpression of *MYC* in the PB2 and PB3 lineages did not noticeably further exacerbate CIN. Note, however, that these cells may have spontaneously increased expression of MYC target genes prior to transduction with the *MYC* lentivirus (Fig. S5). Thus, it is possible that overexpression of *MYC* targets is contributing to the CIN phenotype in the PB2 and PB3 lineages. Whether *MYC* overexpression exacerbates CIN in a *TP53*-mutant only background will require scWGS analysis of P1–3M.

### *TP53* loss initiates extensive transcriptional rewiring

The observation that *TP53* mutant cells accumulate aneuploidies was surprising considering the longstanding observation that p53-null HCT116 cells remain diploid (Bunz et al., 2002; Thompson and Compton, 2010). Indeed, we also found that CRISPR-generated *TP53^-/-^* HCT116 cells do not develop aneuploidies (Simões-Sousa et al., 2018). While *TP53* loss in HCT116 and RPE-1 cells can facilitate tolerance of abnormal karyotypes, p53-activation in response to aneuploidy is not consistent and is context dependent (Santaguida et al., 2017; Simões-Sousa et al., 2018; Soto et al., 2017; Thompson and Compton, 2010). Moreover, it should be noted that such aneuploidy tolerance studies utilised experimental induction of chromosome mis-segregation in cells lacking p53. However, the emergence of aneuploid clones with *TP53* loss has been observed in untreated mammary epithelial and RPE-1 cells (Kok et al., 2020; Salehi et al., 2020; Soto et al., 2017). In addition, multiple cellular processes were deregulated in response to p53 inactivation in transformed murine embryonic fibroblasts, including ploidy control (Valente et al., 2020). Therefore, the fact that *TP53* mutant FNE1 cells accumulate aneuploidies without exposure to exogenous replication stress or mitotic perturbation suggests that, in this context, p53 loss is also sufficient to initiate CIN. To explore potential underlying mechanisms, we performed global transcriptomics, analysing the panel of 18 derivatives by RNAseq. Parental FNE1, P1, P1E and P1M were analysed in triplicate, totalling 27 samples.

A principal component analysis (PCA) yielded four clusters, with cluster 1 comprised of the three parental FNE1 samples (Fig. 6A). Cluster 2 is dominated by the three independent *TP53* mutants, P1–3, and their ‘*empty’* vector derivatives P1–3E, thus reflecting gene expression changes induced by *TP53* loss. Cluster 3 contained the PB2 and PB3 lineages, reflecting the effect of *BRCA1* loss in the *TP53*-mutant background. Cluster 4 contained P1–3M and thus reflects gene expression changes induced by *MYC* overexpression on the *TP53*-mutant background. Note that PB1, and its empty vector derivative PB1E, are in cluster 2, rather than the *BRCA1*-deficient cluster 3. Likewise, PB1M is in cluster 4 with P1–3M. However, as described above, the PB1 lineage may not be fully BRCA1-deficient due to alternative splicing of exon 11. Note also that while overexpression of *MYC* had a marked effect on P1–3 and PB1 cells, it had little effect on the PB2 and PB3 cells. However, again, as described above, these cells appear to have spontaneously upregulated expression of *MYC* target genes (Fig. S5), explaining why ectopic *MYC* had little additional effect. Based on these observations, we conclude that *TP53* mutation alone results in profound transcriptional rewiring, which is further amplified by either elevated MYC activity or BRCA1-loss, in the latter case spontaneous MYC upregulation and MYC-independent enrichment of target genes were observed.

**Figure 6:**
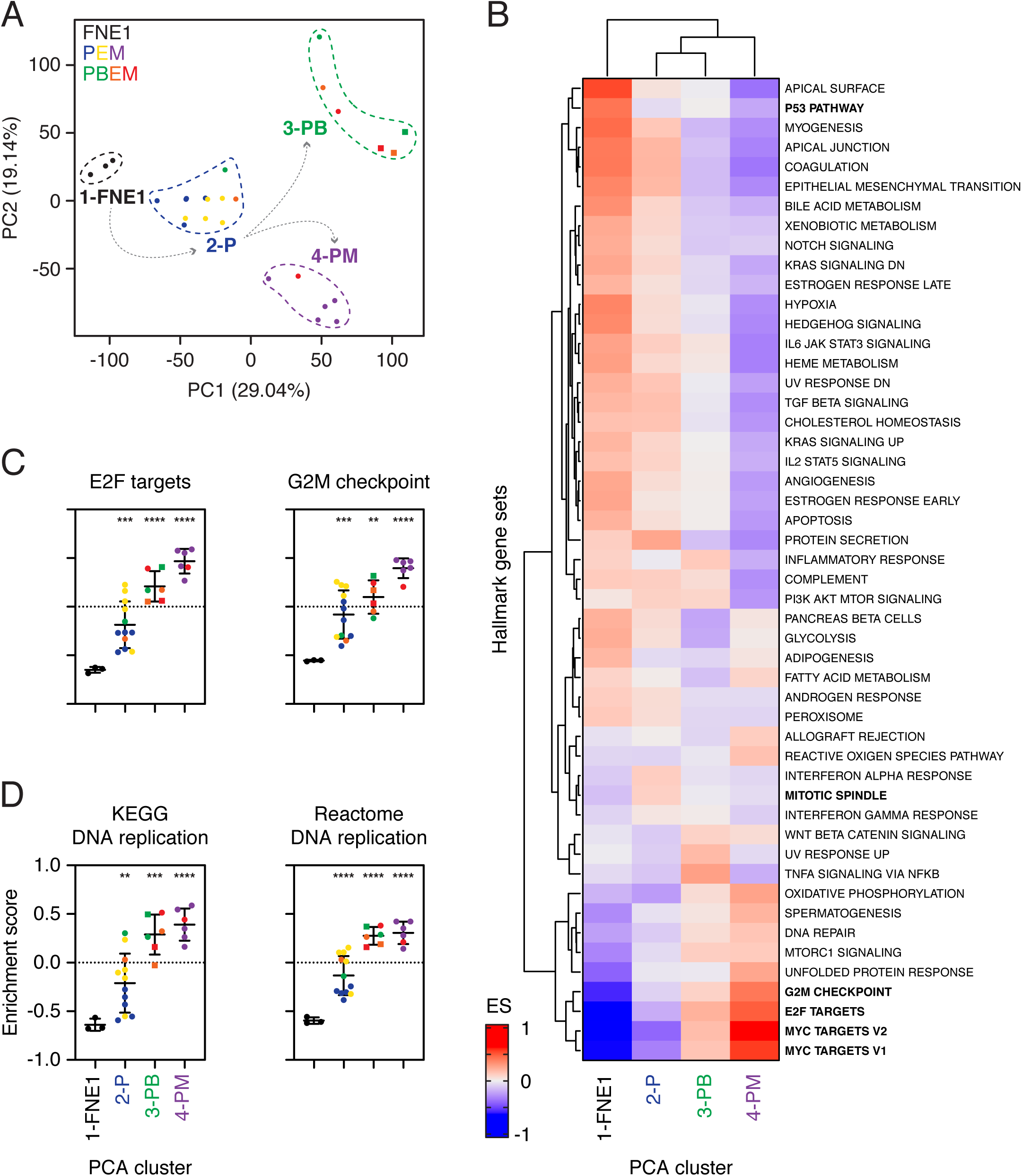
Transcriptome Profiling Reveals Cell Cycle Deregulation Upon p53 Loss. **A** Principal component analysis (PCA) of 27 cell lines analysed by RNA sequencing separates parental FNE1 samples from mutant subclones and BRCA1-deficient subclones from those with fully or partially functioning BRCA1. Indicated colours correspond to sample genotype. Dotted lines capture four clusters defined by similarity of transcriptomes that broadly follow sample genotype with the exception of PB1 and PB1E/M (see text). Samples derived from the PB3 lineage are depicted as squares. Percent variance of principle components 1 (PC1) and 2 (PC2) are indicated in parenthesis along axes. See corresponding Table S3 for input data. **B** Gene set variation analysis (GSVA) was performed on samples grouped according to each of the four distinct PCA clusters and the mean was used to perform unsupervised hierarchical clustering. The 50 Hallmark gene sets are indicated, and the enrichment score (ES) is depicted in blue or red for negative or positive enrichment, respectively. See also Figure S4 and Table S4. **C–D** Results from two representative Hallmark gene sets from B, and the DNA replication gene sets from the KEGG and Reactome collections are shown. Samples were grouped based on PCA cluster allocation and the colour of individual data points corresponds to sample genotype as in A. Samples derived from the PB3 lineage are depicted as squares. For cluster 1 (FNE1): n=3 samples; cluster 2 (P): n=12; and clusters 3 and 4 (PB and PM): n=6. Horizontal bar and error bars indicate mean and standard deviation, respectively. Asterisks depict adj. p-value between indicated groups compared with cluster 1 (FNE1) by Brown-Forsythe and Welsh ANOVA where * adj. p-value ≤ 0.05, ** adj. p-value ≤ 0.005, *** adj. p-value ≤ 0.0005, **** adj. p-value < 0.0001. See Figure S5 and Table S5. P=*TP53*-mutant; B=*BRCA1*-mutant; E=empty vector lentivirus; M=*MYC*-overexpressing lentivirus.

### *TP53* loss deregulates cell cycle gene expression programmes

To determine how *TP53* and *BRCA1* loss and *MYC* overexpression deregulate the transcriptome in FNE1 cells, we performed gene set variation analysis (GSVA) using the Hallmark gene set collection, an approach that allows comparisons across multiple sample groups (Hänzelmann et al., 2013). Unsupervised hierarchical clustering of the 27 samples resulted in a similar separation as the PCA, with parental FNE1 (cluster 1) and the *TP53* mutants (cluster 2) forming one clade (Fig. S4). The *TP53* mutants overexpressing *MYC* (cluster 4) formed a separate clade, while the *BRCA1*-deficient lineages PB2 and PB3 (cluster 3) formed a further two clades. Next, we grouped the various cell lines into the four PCA clusters and interrogated specific gene sets. Consistent with p53 proficiency, the p53 pathway gene set was positively enriched in the parental FNE1 group (cluster 1) versus the *TP53*-mutant lineages (clusters 2–4, Fig. 6B, S5). *MYC* target gene sets V1 and V2 were most highly positively enriched in cluster 4, i.e., the *TP53*-mutant samples overexpressing *MYC* (Fig. 6B, S5). *MYC* targets were also enriched in the PB2 and PB3 lineages (cluster 3), despite only two of the six lines harbouring ectopic *MYC*, demonstrating spontaneous upregulation of *MYC* targets in PB2 and PB3. E2F targets, G2/M checkpoint and mitotic spindle gene sets also stand out; in all three cases, parental FNE1 cells (cluster 1) display negative enrichment, which suggests attenuation of these genes’ expression in a p53-proficient background. Consequently, as genetic manipulations are introduced, the enrichment score progressively increases (clusters 2–4; Fig. 6C, S5). Importantly, because cluster 2 cells showed significant increases in enrichment score versus parental FNE1 cells for E2F targets, MYC targets, G2/M checkpoint and mitotic spindle gene sets, these observations indicate that loss of p53 is sufficient to deregulate multiple aspects of cell cycle control (Fig. 6C, S5). Conversely, this reveals a surprising role for wildtype p53; in the absence of cellular stresses predicted to hyper-stabilize p53, basal levels of p53 appear to be, either directly or indirectly, repressing expression of genes governing a range of cell cycle controls.

### *TP53* loss deregulates expression profiles of DNA replication genes

As replication stress is an established CIN driver (Burrell et al., 2013; Tamura et al., 2020), we next asked whether evidence of replication stress manifested in the RNAseq data. Indeed, upregulation of DNA replication genes is an established mechanism to tolerate replication stress (Bianco et al., 2019). However, because the Hallmark collection does not contain a DNA replication gene set, we analysed the DNA replication gene sets from the Kyoto Encyclopedia of Genes and Genomes (KEGG) and Reactome collections. GSVA revealed that the DNA replication gene sets showed significant increases in enrichment score versus parental FNE1 cells (Fig. 6D). While the enrichment score remains negative for the *TP53*-mutants (cluster 2), it is significantly increased compared with parental FNE1 cells, indicating that p53 loss is perhaps sufficient to induce replication stress.

Taken together, our observations indicate that *TP53* mutation is sufficient to deregulate multiple cell cycle gene expression programmes and trigger transcriptional alterations consistent with a response to replication stress, and that these changes are exacerbated by mutation of *BRCA1* and overexpression of *MYC*. Coupled with the ploidy and karyotype analysis, these observations provide a plausible mechanism by which *TP53* loss is sufficient to initiate CIN in FNE1 cells.

### p53-deficient mouse fallopian tube organoids display cell cycle deregulation

Our finding that *TP53* loss is sufficient to deregulate gene expression programmes governing cell cycle progression, DNA replication and mitosis was surprising. Therefore, we asked whether data from an independent model system supported our observation. Recently, a series of mouse fallopian tube organoids have been developed harbouring conditional alleles designed to inactivate *Trp53* and express an SV40 Large T antigen, which in turn suppresses Rb1 function. Utilising the publicly available RNAseq data, we analysed differentially expressed genes and performed GSEA analysis. PCA shows that the wildtype and mutant organoids form two distinct clusters, indicating divergent gene expression profiles (Fig. S6A), and unsupervised hierarchical clustering analysing E2F, G2/M and mitotic spindle-related genes clearly separated wildtype from mutant (Fig. S6B). Finally, we correlated the normalized enrichment scores for various gene sets in our human FNE1-derived *TP53*-deficient P cells with the mouse organoid samples. This showed that MYC targets, E2F targets, G2/M checkpoint genes and mitotic spindle genes were all positively correlated in both samples. Thus, although the mouse organoids are deficient for both p53 and Rb1 function, the gene expression changes are mirrored in human FNE1 cells harbouring mutant *TP53*, further supporting our notion that p53 loss in human FNE1 cells is sufficient to drive profound transcriptional deregulation of cell cycle regulators.

### *TP53* loss confers tolerance to pharmacologically induced mitotic perturbation

Our observations show that in FNE1 cells, *TP53* mutation is sufficient to induce CIN, and that this is accompanied by deregulation of gene expression networks required to maintain chromosomal stability. As gene expression profiling only indirectly reflects cell function, we asked whether *TP53* mutation does indeed modulate the functionality of chromosome stability pathways. To do this, we challenged parental FNE1 cells and *TP53-*deficient P1 cells with GSK923295, an inhibitor of the mitotic kinesin CENP-E (henceforth CENP-Ei), and analysed the effects by time-lapse microscopy, using cell confluency as a proxy for proliferation. Note that pharmacological inhibition of CENP-E prevents congression of a small number of chromosomes, thus preventing satisfaction of the spindle assembly checkpoint (SAC), in turn inducing a mitotic arrest. Eventually, ‘SAC exhaustion’ results in anaphase onset and mitotic exit in the presence of polar chromosomes, leading to aneuploidy (Bennett et al., 2015; Wood et al., 2010).

In the absence of inhibitor, both populations proliferated and then reached a confluency plateau after 48 hours (Fig. 7A). Upon exposure to CENP-Ei, both parental FNE1 and P1 cells underwent mitotic arrest, evidenced by a static confluence during the first 12 hours and an increase in mitotic index (Fig. 7A,B). They eventually divided and flattened out, resulting in a confluence increase. Parental FNE1 cells failed to divide again, yielding a long second plateau and progressive decrease in mitotic index. By contrast, *TP53-*mutant P1 cells entered and exited a second mitosis, indicated by a short second plateau followed by sustained confluency increase and consistently increased mitotic index (Fig. 7A,B). To confirm this, we performed cell fate profiling, analysing 25 individual cell divisions and tracking the fate of the daughters. In the absence of CENP-Ei, cells in both populations completed multiple rounds of cell division (Fig. 7C). Upon exposure to CENPEi, both parental FNE1 and P1 cells underwent prolonged mitotic delays (Fig. 7C, compare the length of black bars), but, following eventual exit, while the parental FNE1 cells were then blocked in the subsequent interphase, the vast majority of the p53-deficient P1 cells entered second mitoses, indicating continued cell cycle progression.

**Figure 7:**
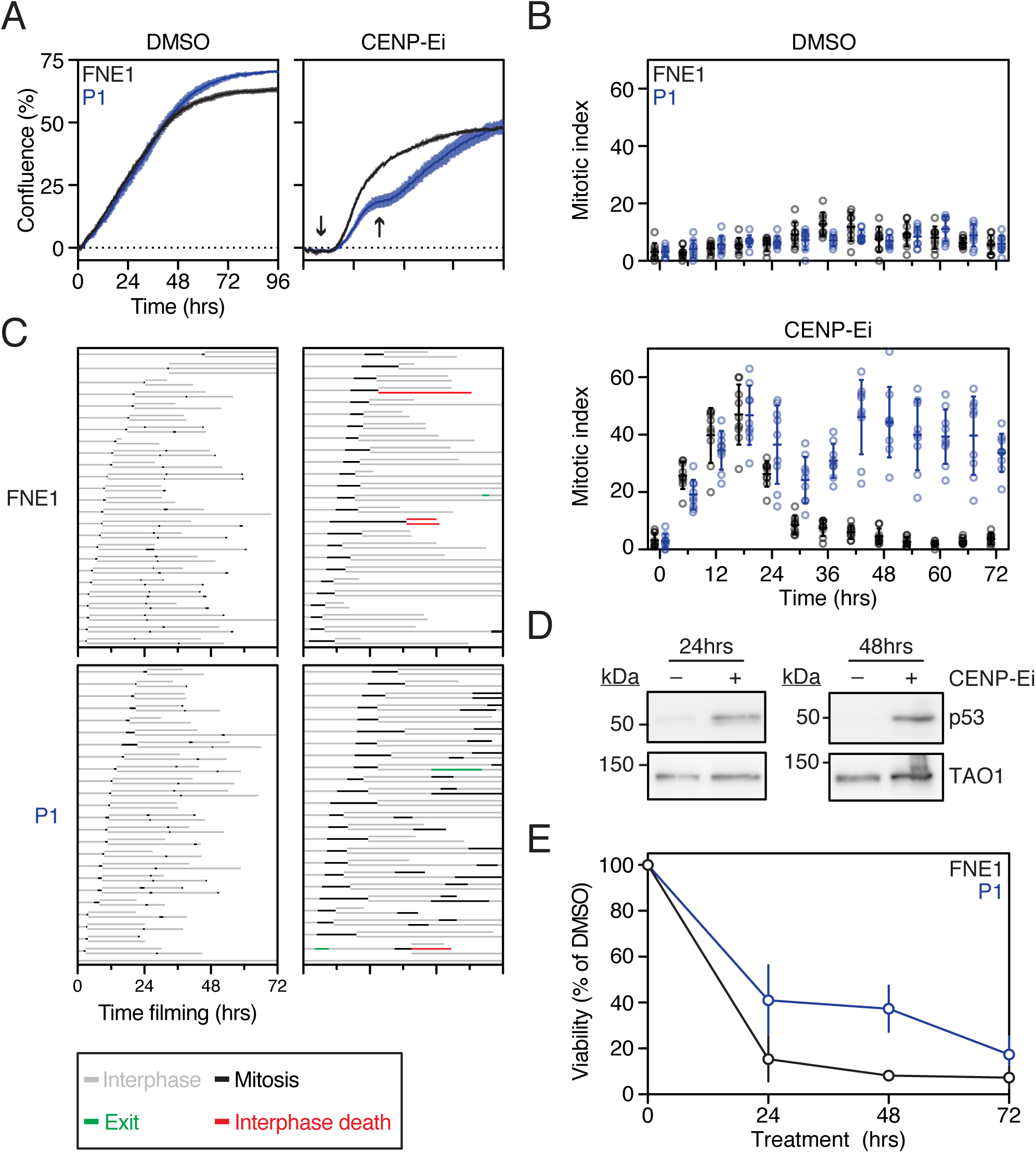
p53 Loss Alone Permits Pharmacologically Induced CIN. **A** Confluence curves of parental FNE1 and *TP53*-mutant (P1) cells in the presence of DMSO (vehicle) or CENP-Ei (GSK923295). Confluence was normalized to T_0_ by subtraction. Arrow indicates mitotic arrest. Representative results from three technical replicates of at least three independent experiments are shown. Error bars represent standard deviation. **B** Mitotic index was profiled in parental FNE1 and P1 cells in the presence of DMSO or CENP-Ei at indicated time points. Results shown are from three fields of view from three technical replicates shown in A. **C** Cell fate profiling by time-lapse microscopy of parental FNE1 and P1 cells in the presence of DMSO or CENP-Ei. 25 cells and both daughters of the first mitosis were profiled per condition. **D** Immunoblot of p53 expression in parental FNE1 cells treated with DMSO or CENP-Ei for 24 and 48 hours. TAO1 serves as loading control. **E** Crystal violet-based viability assay of parental FNE1 and P1 cells treated with DMSO or CENP-Ei for indicated time period followed by drug washout. Experiment was concluded 14 days after drug addition and viability was normalized to DMSO-treated cells. Two independent experiments are shown for the 24- and 72-hour washouts and three for 0- and 48-hour washouts. Error bars represent standard deviation.

Consistent with the interphase block, p53 was stabilised in parental FNE1 cells (Fig. 7D) and longer-term viability was diminished (Fig. 7E). Thus, we conclude that loss of *TP53* in FNE1 cells is sufficient to compromise the post-mitotic cell cycle blocks that would normally prevent proliferation of aneuploid daughter cells following a prolonged mitosis and chromosome mis-segregation event. While we have not analysed the effect of p53 loss on replication stress and G2/M checkpoint controls directly, these observations are consistent with the notion that *TP53* disruption is sufficient to compromise cell biological processes that would otherwise function to minimise CIN.

## DISCUSSION

HGSOC is characterized by ubiquitous mutations in *TP53* and high levels of aneuploidy as a consequence of CIN (Cancer Genome Atlas Research, 2011; Ciriello et al., 2013). However, a genetic basis for CIN in HGSOC remains elusive. In this study, we set out to investigate whether the genetic alterations commonly observed in HGSOC are sufficient to drive CIN, in particular in the HRD group characterized by *BRCA1/2* mutation and *MYC* amplification (Wang et al., 2017). As HGSOC predominately originates from the fallopian tube, we generated a panel of CRISPR/Cas9-mutant, fallopian tube-derived sub-clones based on the *hTERT*-immortalized, non-transformed cell line FNE1 (Labidi-Galy et al., 2017; Merritt et al., 2013). We first showed that FNE1 cells mount a robust p53 response indicating pathway proficiency, in contrast to other model cell lines which rely on p53 suppression for immortalization (Fig. S1A,B) (Karst and Drapkin, 2012; Karst et al., 2011; Nakamura et al., 2018). Importantly, parental FNE1 p53 proficiency allowed us to directly test the impact of p53 loss of function alone, and in combination with BRCA1 deficiency and MYC overexpression, in an isogenic model system. Using this system, we find that p53 loss alone is sufficient to cause aneuploidy in FNE1 cells, which is exacerbated in the absence of functional BRCA1. Analysing the transcriptome revealed that cell cycle de-regulation was apparent in *TP53* single mutants and amplified in *TP53*/*MYC* double mutants. The most highly enriched gene sets compared with the parental FNE1 cells were G2/M checkpoint, E2F targets, DNA replication and mitotic spindle, which were enriched in cells deficient for p53 alone and progressively more enriched with additional genetic manipulations. These findings, which were consistent with publicly available data from mutant mouse fallopian tube organoids (Fig. S6) (Zhang et al., 2019), therefore indicate that p53 loss alone results in transcriptional changes that can deregulate the cell cycle and promote low-level CIN. Since truncating mutations that lead to a loss-of-function only account for 35% of HGSOC (Cancer Genome Atlas Research, 2011), future work will require to look into other, missense and potential gain-of-function, *TP53* mutations in this context.

*TP53* mutations have been firmly established as early and ubiquitous events in HGSOC development. However, the implications of *TP53* mutation on fallopian tube epithelial cells remain poorly understood and have thus been highlighted as key to understanding the development of HGSOC (Bowtell et al., 2015). Although p53 has been established as suppressor of proliferation in response to aneuploidy, mutations in *TP53* correlate consistently and most strongly with aneuploidy and WGD in multiple tumour types (Bielski et al., 2018; Ciriello et al., 2013; Davoli et al., 2017; Taylor et al., 2018; Thompson and Compton, 2010; Zack et al., 2013). While evaluation of fallopian tube-derived models with suppressed p53 has previously suggested that additional p53-independent mechanisms act as barriers to proliferation of aneuploid cells, the same study found increased potential of transformation with p53 suppression in combination with pharmacologically induced aneuploidy in soft agar assays (Chui et al., 2019). Conflicting observations have also been reported regarding the relationship between p53 loss and the emergence of aneuploidy in studies utilizing colorectal cancer cell lines (Bunz et al., 2002; Simões-Sousa et al., 2018). Indeed, we observed an increase in structural and numerical aneuploidy by scWGS when comparing parental FNE1 with p53-deficient P1 cells. Although the magnitude of this change is moderate quantitatively, on a qualitative level it is evident that P1 cells harbour more whole chromosome or chromosome arm aneuploidies than parental FNE1 cells from two different passages (Fig. 5). Therefore, mounting evidence from us and others suggests that p53 loss alone may be sufficient to induce low levels of CIN, permitting cells to explore karyotypic heterogeneity. However, the importance of environmental factors such as O_2_ levels has only recently been brought to light which might impact both chromosome segregation and the processes preceding mitosis as well as the selection of explorable karyotypes. It is conceivable that growth conditions at atmospheric O_2_ levels may previously have masked the emergence of aneuploidy as euploid cells would outcompete aneuploid cells more rapidly than under normoxic or hypoxic conditions (Rutledge et al., 2016).

The development of isogenic, *bona fide* mutant cell lines allowed us to study mitotic perturbations side-by-side in p53-proficient and -deficient cells. HGSOC is appreciated as one of the most chromosomally unstable cancer entities based on *in silico* analyses of cancer genomes, which were backed up by cell biological studies of mitosis in HGSOC models (Nelson et al., 2020; Tamura et al., 2020). Primary cultures established from HGSOC patients’ ascitic fluid can take more than six hours to complete mitosis in extreme cases, and up to 24 hours in select examples of individual cells (Nelson et al., 2020). This dramatically increased mitotic duration compared with non-transformed cells has been shown to be limited in a p53-dependent manner termed the ‘mitotic timer’. Indeed, knockout of *TP53* and its upstream regulators in this specific context, *USP28* and *53BP1*, rescued growth arrest following prolonged mitosis of up to six hours (Lambrus et al., 2016). Inhibiting the mitotic kinesin CENP-E pharmacologically, we could achieve a comparable increase in mitotic duration and were able to show that p53 was stabilized in response to CENP-E inhibition. Furthermore, we show that P1 cells tolerate this stress better than parental FNE1 cells in short-term as well as long-term assays (Fig. 7). Thus, we show that p53 loss precipitates low levels of CIN and also partially rescues viability upon mitotic delay and chromosome mis-segregation; this dual- or potentially multi-functionality of p53 provides an explanation as to why one of the most chromosomally unstable tumour entities is characterized by ubiquitous *TP53* mutations.

Beyond mutations in *TP53*, mutations in *BRCA1*/*2* are the second most common mutation in HGSOC (12% of cases each). In genetically engineered mouse models of mammary epithelial cancer, deletion of exon 11 of *BRCA1* was shown to cause functional G2/M checkpoint disruption and tumorigenesis (Weaver et al., 2002; Xu et al., 1999). Based on these two observations, and the fact that human BRCA1-deficient fallopian tube-derived cell line models are lacking, we mutated *BRCA1* to create a model of more pronounced CIN and HRD. We found that our three cell lines deficient in full length BRCA1 are distinct from one another; based on the analysis of gene expression profiles by PCA and GSVA, PB1 clusters with P cells and PB2 and PB3 each form independent clusters. This distinction likely reflects biological heterogeneity following *BRCA1* mutagenesis that led to exacerbation of CIN. Indeed, PB1 cells are largely 2c, while PB2 cells harbour a 2c and 4c population and PB3 cells are 4c. Interrogation of our RNAseq data on the nucleotide level found that PB2 and PB3 have an identical exon 3 mutation, however, PB1 cells express a splice variant of exon 11 as a consequence of a mutation in the same exon, which is known to diminish PARPi sensitivity versus other BRCA1-mutants (Wang et al., 2016). Our findings are in agreement with this *BRCA1* variant having some functionality, as we find that, despite the absence of full-length *BRCA1*, its retained expression might be sufficient to protect against aneuploidy. As flow cytometric and miFISH evidence suggested aneuploidy, PB2 and PB3 were subjected to scWGS and indeed the extent of copy number heterogeneity observed exceeded that of P1 cells. Interestingly, we observed a propensity for WGD in both PB2 and PB3, despite *BRCA1* mutations not being reported to correlate with whole genome doubling (Bielski et al., 2018). This could reflect an *in vitro* selection pressure permitting the detection of 4c PB2 and PB3 cells in our system. Nevertheless, we conclude that the combination of p53- and BRCA1-deficiency can drive CIN in a context-dependent manner, where low levels of BRCA1 activity such as observed in PB1 remain protective.

Several non-genetic causes of CIN such as increased microtubule assembly rates, centrosome amplification and replication stress have been identified in colorectal cancer and HGSOC cell lines (Bastians, 2015; Tamura et al., 2020). To try and decipher the causes of CIN in our mutant subclones we turned to analysis of transcriptomics, which enabled us to take an agnostic, genome-wide approach. We observed that loss of p53 alone resulted in an enrichment of gene sets comprised of genes regulating the cell cycle and DNA replication. We suggest that this effect is a consequence of the downregulation of canonical p53-targets such as *MDM2* and *CDKN1A*, which encodes the CDK inhibitor p21 (Fig. 4C). p21 plays an important role in suppressing S-phase entry by negatively regulating cyclin E and CDK2. The absence of this negative regulation thus permits cyclin E and CDK2 to hyperphosphorylate RB1 more rapidly, which results in de-sequestration of E2F, a key transcription factor controlling S-phase entry (Sullivan et al., 2018). Indeed, the E2F targets gene set is significantly less negatively enriched in P samples than in parental FNE1 samples (Fig. 6C). To contextualize, p21 has been shown to protect cells from CIN. In a genetically engineered mouse model of p53 separation-of-function, which was apoptosis-deficient but partially functional to suppress cell cycle progression, deletion of p21 led to an increase in CIN (Barboza et al., 2006). Moreover, three of the four sample groups showed significantly different and more positive enrichment scores in cell cycle related gene sets compared with parental FNE1 cells.

With the exception of the mitotic spindle gene set, overexpression of *MYC* consistently amplified the already observed enrichment in p53-deficient P samples, likely reflecting MYC’s role as transcriptional amplifier (Lin et al., 2012; Nie et al., 2020; Nie et al., 2012). This held true also for the negative enrichment of the p53 pathway gene set where P samples displayed an already negative enrichment score that was even more negative in the PM samples (Fig. S5). In contrast to P samples, *MYC* overexpression did not seem to have the same impact on the transcriptome in PB2 and PB3 as it did in PM samples (Fig. 6A). In fact, PB2 and PB3 showed more positive enrichment of MYC targets V1 and V2 than P samples even without *MYC* overexpression; this is consistent, at least in PB3 samples, with higher endogenous *MYC* transcript levels (Table 1). Interestingly, the PB2M sample reaches the highest enrichment score of the PB2 lineage suggesting that ectopic MYC is active in this sample, but perhaps to a lesser extent than in PM samples. Consistent with our findings, proteogenomic analyses of HGSOC had suggested a causal role for the deregulation of mitotic and DNA replication genes in the high levels of CIN observed in this disease, however, the causes for this deregulation could not be definitively dissected in patient samples (McDermott et al., 2020). Taking these data into account, we suggest that CIN is caused by the cumulative changes in cell cycle regulators’ expression, rather than a single causative gene, as a consequence of, e.g., loss of p53-signalling through its downstream effector p21, which promotes transcriptional programs of cell cycle progression. Future work should focus on genetic add-back experiments of down-regulated *CDKN1A* (encodes p21) to investigate if this rescues the observed deregulated expression of cell cycle genes and low-level CIN.

In summary, we provide evidence, based on a novel human, fallopian tube-derived cell line panel that p53 loss leads to transcriptomic deregulation of cell cycle regulators, which is amplified by the overexpression of the oncogene *MYC*. We propose that the sum of these transcriptional changes causes CIN in HGSOC and show that P1 cells display low levels of aneuploidy. Furthermore, we show that additional genetic manipulation of *BRCA1* exacerbated both the enrichment of cell cycle regulators and aneuploidy. Finally, p53 loss increased tolerance of pharmacological perturbation of mitosis using an inhibitor of CENPE, further supporting its potential role in the development of CIN in HGSOC. Our data point to the dual- or multi-functional role of p53 whereby its loss precipitates CIN by causing cell cycle and DNA replication deregulation while simultaneously also promoting the survival of aneuploid cells that experienced those stresses in the previous cell cycle.

## MATERIALS AND METHODS

Details of critical commercial reagents and kits, drugs, antibodies, recombinant DNA, oligonucleotides, FISH probes and software are contained in Table S1.

### Cell culture

FNE1 cells (a kind gift from Dr Tan A. Ince) were cultured in WIT-Fo Culture Media (FOMI) at 5% O_2_ and 5% CO_2_ at 37°C, as described previously (Merritt et al., 2013). AAV293T cells (ATCC) were cultured in DMEM supplemented with 10% FBS and 100 U ml^−1^ penicillin-streptomycin, at atmospheric O_2_ and 5% CO_2_ at 37°C. All cell lines were authenticated using the Promega Powerplex 21 System and regularly tested for Mycoplasma either by PCR (both at CRUK Manchester Institute Molecular Biology Core Facility) or the Lonza enzymatic test (Animal Molecular Diagnostics Laboratory at NCI Frederick, MD).

Lentiviruses were produced by co-transfection of AAV293T cells at 5 × 10^4^ cells per well in a 24-well microplate with recombinant DNA at 0.375 µg lentivirus of interest, 0.5 µg psPAX2 and 0.125 µg pMD2.G (both kind gifts from Dr Didier Trono) using the Promega ProFection Mammalian Transfection System kit according to manufacturer instructions. Transfection media was replaced after overnight incubation and lentivirus was harvested every other day for four days. Supernatant containing lentivirus was centrifuged, filtered (0.45 µm) and frozen for storage at −80°C.

CRISPR/Cas9-expressing FNE1 cells were generated by transduction with Dharmacon Edit-R Inducible Lentiviral Cas9 particles followed by selection with blasticidin S at 8 µg ml^−1^. Cas9 expression was assessed by titrating tetracycline and induced using 15 µg ml^−1^ in subsequent experiments. To mutate *TP53*, FNE1 cells expressing inducible Cas9 were transduced with lentiGuide-Puro (a kind gift from Dr Feng Zhang (Sanjana et al., 2014)) containing a guide RNA (gRNA) targeting *TP53* (Table S2) and selected in 0.7 µg ml^−1^ puromycin. Cas9 was then induced for five days before isolation of single-cell clones by limiting dilution (either immediately or following five days further selection in Nutlin-3). Taking P1 cells forward, cells were transduced with six different lentiGuide-Neo (see ‘Molecular Biology’ for details) lentiviruses each containing a unique gRNA targeting *BRCA1* (Table S2). After neomycin selection at 0.8 mg ml^−1^, Cas9 was induced as above before isolation of single-cell derived subclones by limiting dilution. Clones were screened by immunoblotting (see ‘Biochemistry’ for details). Mutations in targeted genes were assessed in the RNA sequencing dataset using Integrative Genomics Viewer (Version 2.8.0) and annotated according to standard practices (Ogino et al., 2007; Robinson et al., 2011). Mutations in *BRCA1* in PB1 and PB2 cells were confirmed using Sanger sequencing. *MYC* overexpressing and cognate ‘E’ cells were generated by transduction with pLenti CMV Hygro DEST or MYC lentiviruses (a kind gift from Drs Eric Campeau and Paul Kaufman (Campeau et al., 2009)) and selection with 25 µg ml^−1^ hygromycin, maintaining a polyclonal cell population. Immunoblotting and RNA sequencing were employed to confirm functionality of *MYC* overexpression. All lentiviral transductions were performed in 4 µg ml^−1^ poly-brene.

Functional deficiency of p53 and BRCA1 in putative clones was confirmed by exploiting the known synthetic-viable and -lethal relationships with Nutlin-3 and PARPi treatment, respectively. Nutlin-3 assays were performed by seeding 30,000 cells (parental FNE1, P1 and P3 transduced with pLVX mCherry-H2B Puro) into Primaria 24-well microplates. The next day, either vehicle (DMSO) or 10 µM Nutlin-3 (Selleck Chem, TX) were added in phenol red-free media and the cells imaged for 96 hours on an IncuCyte® ZOOM (Satorius AG, Germany) time-lapse microscope housed in a low-oxygen incubator (5% O_2_, 5% CO_2_). IncuCyte® ZOOM custom software was used in real-time to measure confluency and red fluorescent object count and for data analysis. Population doubling for each culture was calculated by performing a log_2_ transformation of the fold-change nuclear count from T_0_ and plotted against time. PARPi (Olaparib, Selleck Chem, TX) sensitivity was assessed by seeding 100 cells directly into drug or vehicle containing media in collagen-coated, black 96-well microplates (Greiner Bio-One North America Inc., NC). Media and drug were replenished every three days. On day seven, 30 µl CellTiter-Blue® (Promega, WI) reagent were added to 150 µl of media and incubated for four hours followed by fluorescence signal measurement on a SpectraMax M2 plate reader (Molecular Devices, CA).

Assays studying the response to CENP-E inhibition were performed using GSK923295 (Selleck Chem, TX). For live-cell imaging, 30,000 cells were seeded into Primaria 24-well microtiter plates, allowed to adhere overnight, vehicle or drug (250 nM) were added the next day and imaging on an IncuCyte® ZOOM time-lapse microscope was performed as described above. Cell fate profiles were analysed manually based on exported MPEG-4 videos. Long-term viability assays were performed by seeding 2,000 cells into Primaria 6-well microtiter plates, allowing the cells to adhere overnight and adding vehicle or drug the next day. Drug washout was performed at indicated timepoints and media replenished every 36–48 hours. Experiments were concluded after 14 days, cells were washed, fixed with 1% formaldehyde (in PBS) and stained with crystal violet (0.05% in dH_2_O). Quantitation was achieved by extracting crystal violet with acetic acid and measuring absorbance on a SpectraMax M2 plate reader.

A summary of all cell lines generated is provided in Table 1 and Figure S2A.

### Cell biology

Cells were harvested normally or *in situ*, lysed in sample buffer (0.35 M Tris pH 6.8, 0.1 g/ml sodium dodecyl sulphate, 93 mg/ml dithiothreitol, 30% glycerol, 50 mg/ml bromophenol blue) and boiled for five minutes. Proteins were resolved by SDS-PAGE and electroblotted by wet transfer onto Immobilion-P membranes (Millipore Sigma, MA). Membranes were blocked in 5% milk in TBS-T (50 mM Tris pH 7.6, 150 mM NaCl, 0.1% Tween-20) and incubated with primary antibodies at indicated concentrations (Table S1) overnight at 4°C. Membranes were then washed with TBS-T and incubated with horseradish-peroxidase-conjugated secondary antibodies (Table S1) for two hours at room temperature. After further washes with TBS-T, detection was performed using EZ-ECL Chemiluminescence Substrate (Biological Industries, Israel) or Luminata Forte Western HRP Substrate (Millipore Sigma, MA). Membranes were imaged on Biospectrum 500 (UVP, CA) imaging system.

For p53 immunofluorescence, parental FNE1 cells were seeded onto collagen-coated 19 mm coverslips, incubated overnight and treated with 10 µM Nutlin-3 for 8 hours. Cells were then washed with PBS, fixed (1% formaldehyde in PBS), quenched with glycine, permeabilized with PBS-T (PBS, 0.1% Triton X-100), incubated consecutively with primary (mouse anti-p53, DO-1, Santa Cruz Biotechnology, TX) and secondary (donkey anti-mouse conjugated with Cy3, Jackson ImmunoResearch Laboratories Inc., PA) antibodies for 30 minutes each with a wash step in between (Table S1). Coverslips were then washed with PBS-T, stained with Hoechst 33258 (Millipore Sigma, MA), washed with PBST and mounted onto slides (90% glycerol, 20 mM Tris, pH 9.2). Slides were imaged on an Axioskop2 microscope fitted with a 40× oil immersion objective (both from Zeiss Inc., Germany) and a CoolSNAP HQ camera (Photometrics, AZ) operated by MetaMorph software (Molecular Devices, CA). Image analysis was performed with Adobe Photoshop® CC 2015 (Adobe Systems Inc., CA). Microtiter plates were imaged after addition of PBS on Lionhart FX automated microscope fitted with a 40× objective operated by custom Gen5 (all BioTek, VT) software, which was also utilized for image analysis. RAD51 immunofluorescence was performed as described previously (Callen et al., 2020). Briefly, cells were seeded into a black 96-well microplate (Greiner Bio-One North America Inc., NC) coated with gelatine. Prior to γ-irradiation (5 Gy, ^137^Cs Mark 1 irradiator, JL Shepherd, CA), cells were incubated with 10 µM EdU for 30 minutes. Four hours post-irradiation, cells were pre-extracted (20 mM HEPES, 50 mM NaCl, 3 mM MgCl_2_, 0.3 M sucrose, 0.2% Triton X-100) on ice for 5 minutes to remove soluble nuclear proteins. Pre-extracted samples were fixed (4% paraformaldehyde in PBS), permeabilized (PBS, 0.5% Triton X-100), and incubated with anti-RAD51 antibody (rabbit anti-RAD51, 1:250, Abcam). Detection of RAD51 and EdU was accomplished by incubating samples with Alexa Fluor 568-conjugated secondary antibodies (goat anti-rabbit, Thermo Fisher Scientific, MA) followed by a click-IT reaction as per manufacturer’s instructions (Thermo Fisher Scientific, MA). Finally, DNA was counterstained with DAPI (Thermo Fisher Scientific, MA). Microtiter plates were imaged at 40× magnification on a Lionheart LX automated microscope (BioTek Instruments, Inc.). Quantification of nuclear RAD51 foci was performed using the Gen5 spot analysis software (BioTek Instruments, Inc.).

### Molecular biology

pLenti CMV Hygro DEST (w117-1) was digested with SalI and BamHI (New England BioLabs Inc., MA) according to manufacturer instructions. *MYC* cDNA was PCR-amplified from pcDNA5 FRT/TO CR MYC and cloned into pLenti CMV Hygro DEST, creating pLenti CMV Hygro MYC (Littler et al., 2019). pLVX mCherry N1 (Clonetech Laboratories Inc., CA) was digested with XhoI and BamHI (New England BioLabs Inc., MA) according to manufacturer instructions. H2B cDNA was PCR-amplified from pcDNA5 FRT/TO GFP-H2B and cloned into pLVX mCherry N1, creating pLVX mCherry-H2B Puro (Morrow et al., 2005). Gibson Assembly was utilized to create lentiGuide Neo. Briefly, lentiGuide Puro was PCR-amplified, omitting the puromycin-resistance cassette. Separately, the neomycin-resistance cassette was PCR-amplified from pLXV MYC-mCherry Neo. Fragments were then assembled into lentiGuide Neo using Gibson Assembly Master Mix (New England BioLabs Inc., MA) according to manufacturer instructions. gRNAs were introduced into lentiGuide Puro/Neo by ligating the annealed forward and reverse oligonucleotides into BsmBI-digested target vectors (Sanjana et al., 2014). All recombinant vectors were grown in XL1-Blue competent cells and extracted using QIAprep Spin Miniprep kit (Qiagen, Germany) according to manufacturer instructions. Oligonucleotide sequences are described in Table S2. Recombinant vectors were validated functionally *in vitro* or by Sanger sequencing.

### Molecular cytogenetics

For SKY, cells were cultured as normal and incubated in 100 ng ml^−1^ Colcemid (Roche, MA) for 2 hours prior to harvest. Subsequently, for SKY and miFISH, cells were harvested, swelled in hypotonic buffer (0.075 M KCl) for 30 minutes at 37°C, fixed in methanol/acetic acid (3:1) in three wash steps, dropped onto glass slides and aged for 2 weeks at 37°C. Four probe panels containing five probes each were assembled totalling one centromere probe (CCP10) and 19 gene probes (all custom ordered from CytoTest, MD): *COX2* (1q31.1), *PIK3CA* (3q26.32), *FBXW7* (4q31.3), *CCNB1* (5q13.2), *DBC2* (8p21.3), *MYC* (8q24.21), *CDKN2A* (9p21.3), *PTEN* (10q23.31), *CCND1* (11q13.3), *KRAS* (12p12.1), *RB1* (13.14.2), *CDH1* (16q22.1), *TP53* (17p13.1), *NF1* (17q11.2), *HER2* (17q12), *SMAD4* (18q21.2), *CCNE1* (19q12), *ZNF217* (20q13.2) and *NF2* (22q12.2). Images were taken on an automated fluorescence microscope fitted with a 40× oil immersion objective (BX63, Olympus, Japan), custom optical filters (Chroma, VT) and a motorized stage. Custom software was used for operation and analysis (BioView, Israel). A total of 100 nuclei were analysed per sample for miFISH and 15 metaphases were analysed per sample for SKY. Procedures pertaining to SKY and miFISH hybridization, stripping and rehybridization were as described previously (Heselmeyer-Haddad et al., 2012; Padilla-Nash et al., 2006; Wangsa et al., 2018).

### Next generation sequencing

RNA was extracted from logarithmically growing cells *in situ* using the RNeasy Plus Mini kit (Qiagen, Germany) according to manufacturer instructions. RNA integrity and quality were assessed using a 2200 TapeStation (Agilent Technologies, CA; performed by the CCR Genomics Core, Bethesda, MD). Libraries were prepared using Illumina TruSeq® Stranded mRNA Library Prep (Illumina Inc., CA), pooled and paired-end sequenced on Illumina NovaSeq using an SP flow cell according to manufacturer instructions (Sequencing Facility at NCI Frederick, MD). Samples returned 37 to 51 million pass filter reads with more than 91% of bases above the quality score of Q30.

scWGS was performed on single cells sorted for a 2c (parental FNE1, P1) or 4c (PB3, PB3E, PB3M) genome content (for PB2, PB2E and PB2M 12 cells from each population were included) as described previously (Bakker et al., 2016; Nelson et al., 2020; van den Bos et al., 2016).

### Bioinformatics

For RNA sequencing, sample reads were processed using the CCBR Pipeliner utility (https://github.com/CCBR/Pipeliner). Briefly, reads were trimmed for adapters and low-quality bases using Cutadapt (version 1.18) (http://gensoft.pasteur.fr/docs/cutadapt/1.18) before alignment to the human reference genome (hg38/Dec. 2013/GRCh38) from the UCSC browser and the transcripts annotated using STAR v2.4.2a in 2-pass mode (Dobin et al., 2013; Martin, 2011). Expression levels were quantified using RSEM (version 1.3.0) (Li and Dewey, 2011) with GENCODE annotation version 30 (Harrow et al., 2012). The same approach was used for mouse model data downloaded from Gene Expression Omnibus (GEO, accession number GSE125016), with alignment to the mouse reference genome (mm10).

Raw read counts (expected counts from RSEM) were imported to the NIH Integrated Data Analysis Platform for downstream analysis. Low count genes (counts-per-million [CPM] <0.5), ≥ three samples were filtered prior to the analysis. Counts were normalized to library size as CPM and the voom algorithm (Law et al., 2014) from the Limma R package (version 3.40.6) (Smyth, 2004) was used for quantile normalization (Tables S4 and S7). Batch correction was performed prior to analysis using the ComBat function in the sva package (Johnson et al., 2007). Differentially expressed genes (DEG) using Limma and pre-ranked gene set enrichment analysis (GSEA) were computed between each genotype using the molecular signatures database (Liberzon et al., 2011; Subramanian et al., 2005). And gene set variation analysis (GSVA) was performed using the GSVA package (Hänzelmann et al., 2013). Genes or gene sets with an adjusted p-value ≤0.05 were considered statistically significant. Preparation of heatmaps was performed in R Studio (Subramanian et al., 2005).

Analysis of copy-number changes based on scWGS was executed according to previous reports (Bakker et al., 2016; Nelson et al., 2020; van den Bos et al., 2016).

### Quantification and statistical analysis

Prism 8 (GraphPad, CA) was used to generate graphs and perform statistical analyses. RStudio (R Project for Statistical Computing) was used to perform sequencing analyses and generate heatmaps (R packages Complex Heatmaps and AneuFinder) and volcano plots (R package Enhanced Volcano).

## ACKNOWLEDGEMENTS

We thank members of the Ried and Taylor laboratories for advice. We thank the following facilities: Molecular Biology Core, Imaging and Flow Cytometry Core (both Cancer Research UK Manchester Institute), OSTR Sequencing, Animal Diagnostics Laboratory (both NCI at Frederick, MD), CCR Genomics Core and Laboratory of Genome Integrity Flow Cytometry Core (both NCI at Bethesda, MD).

## COMPETING INTERESTS

We report no competing interests.

## FUNDING

DB was supported by a Wellcome Trust-NIH PhD Studentship (200932/Z/16/Z) and the NIH-OxCam program. DH was supported by a Mildred Scheel postdoctoral fellowship from the Deutsche Krebshilfe. This study was in part supported by the intramural research program of the National Institutes of Health, the National Cancer Institute. This study was also in part supported by a Cancer Research UK Program grant to SST (C1422/A19842) and a Cancer Research UK Centre Award (C5759/A25254).

## DATA AVAILABILITY

Next generation sequencing data will be made available without restriction through GEO or the EMBL-EBI’s repository upon publication in accordance with the journal’s publishing policy.

## AUTHOR CONTRIBUTIONS STATEMENT

All experiments and analyses were performed by DB except for the following: Fig. S1C and Fig. 5 were performed by RW, DS and supervised by FF; Fig. S1D was performed by DW; Fig. 2D was performed in part by DZ and supervised by AN; Fig. 2E, 3A,C,D, 6, S2B, S4, S5 and S6 were performed with help from TJM and supervised by MC. All other authors provided reagents and/or technical support. TR and SST provided additional funding and supervision. DB interpreted the data and wrote this manuscript, both with support from JM, TR and SST. All authors read the manuscript and provided feedback.

---

**Figure S1:**
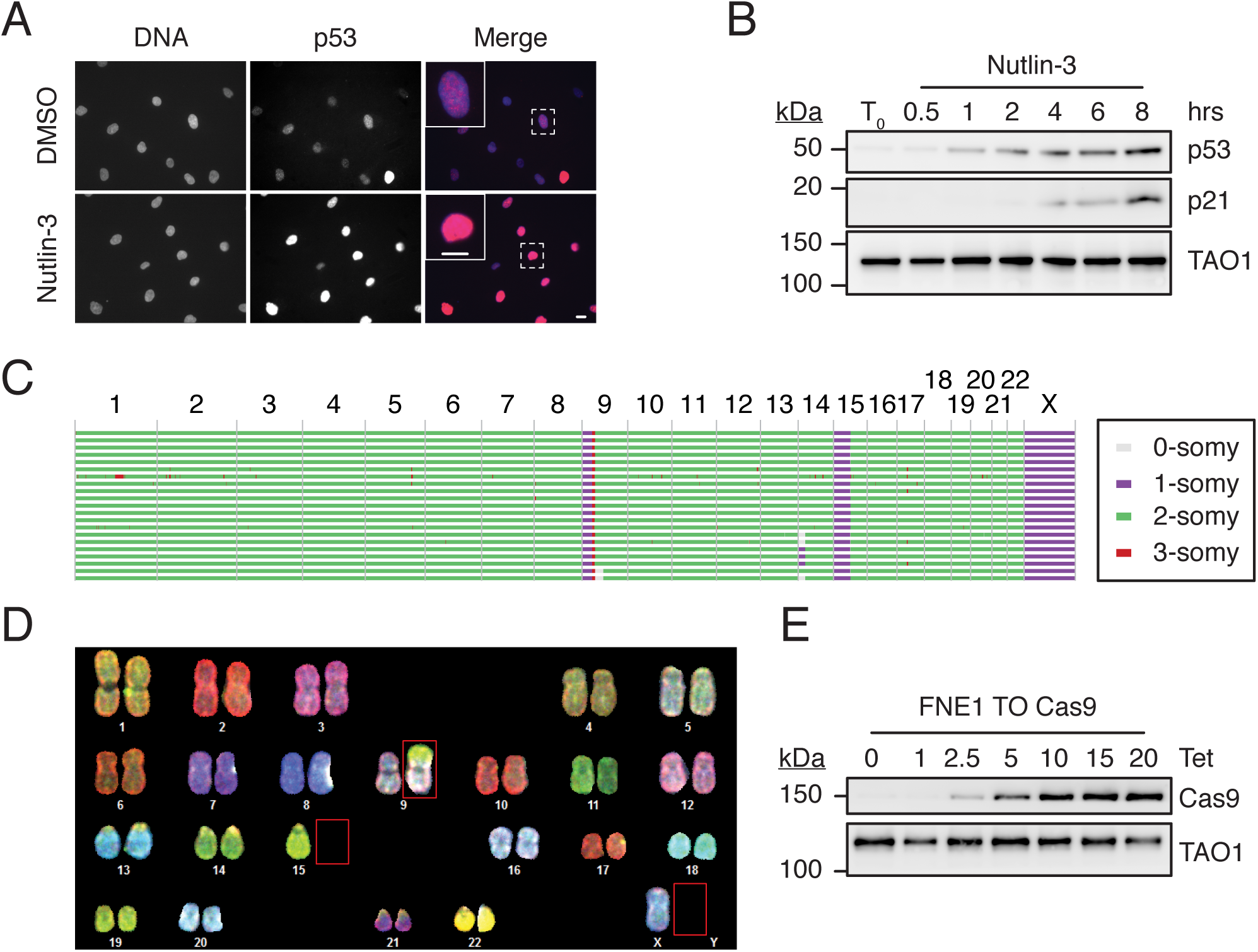
FNE1 Characterization. **A** Immunofluorescence imaging of DMSO (vehicle) and Nutlin-3-treated parental FNE1 cells shows stabilization of p53 in response to Nutlin-3. Representative images from one of three experiments. Scale bars, 10 μm **B** Immunoblot of cells treated with Nutlin-3 over a time course of 8 hours to analyse p53 and p21 expression. TAO1 serves as loading control. **C** Shallow-depth whole-genome sequencing analysis of copy number aberrations in single parental FNE1 cells (rows) where columns reflect chromosomes 1–22 and X. Colour indicates detected copy number level (box). **D** Spectral karyotyping image of a representative metaphase spread shows a near-diploid genome with loss of chromosomes 15 and X and translocation between 9p and 15q (red boxes). **E** Immunoblot of tet-inducible Cas9 expression in parental FNE1 cells after transduction with Edit-R Inducible Lentiviral Cas9 and selection. Subsequent experiments utilized 15 µg ml^−1^ tet for Cas9 induction. TAO1 serves as loading control. Tet= μg ml^−1^ tetracycline.

**Figure S2:**
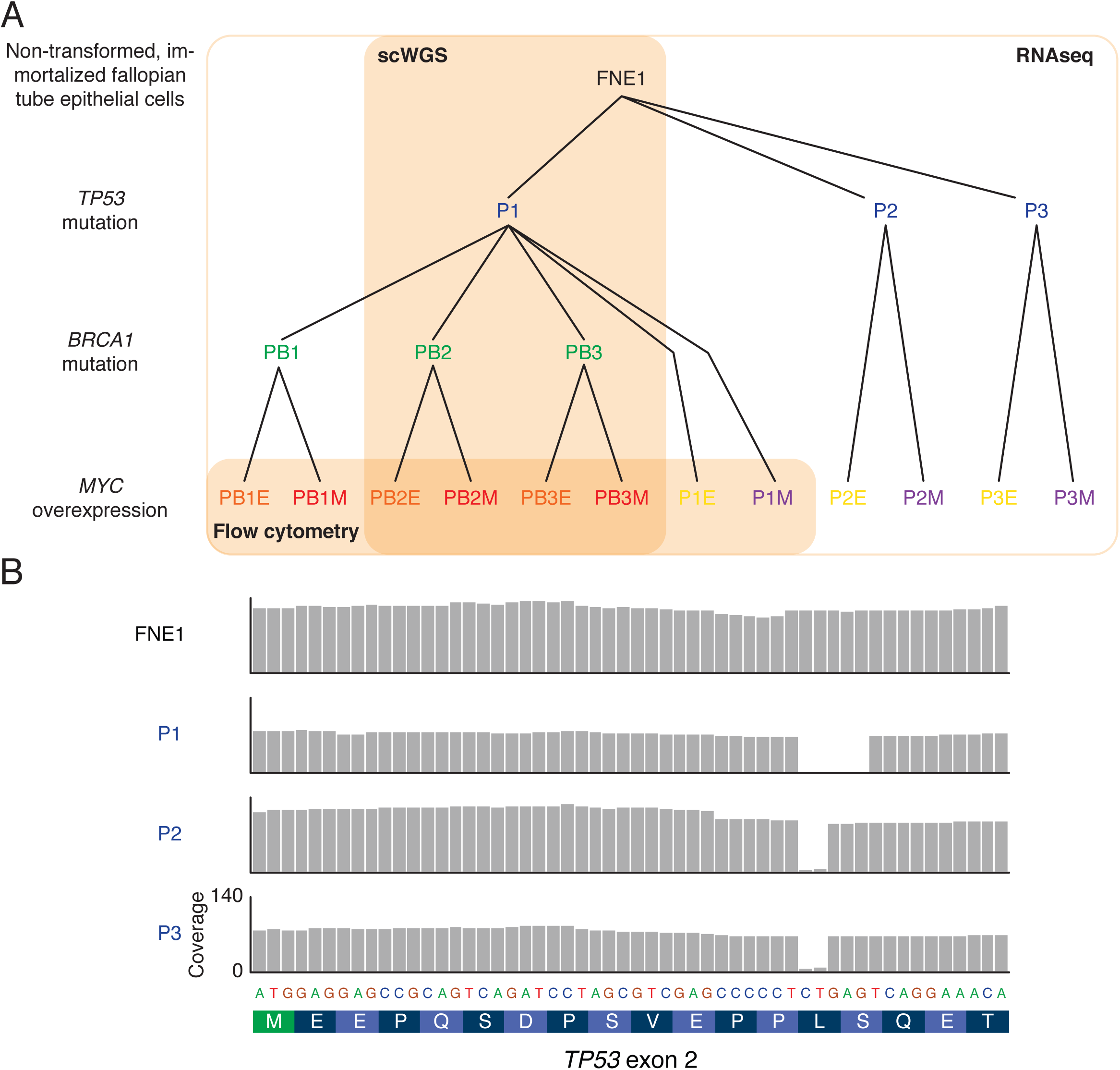
Pedigree of Mutant Subclones and *TP53* Locus Mutation. **A** Pedigree of FNE1 cells and sequentially CRISPR/Cas9-mediated genome-engineered subclones with introduction of MYC overexpression or empty lentiviral construct. **B** Coverage of RNA sequencing reads of *TP53* exon 2 in indicated subclones. Deletion of 2–5 nucleotides in the three mutagenized subclones is shown, resulting in a downstream premature termination codon. P=*TP53*-mutant; B=*BRCA1*-mutant; E=empty vector lentivirus; M=*MYC*-overexpressing lentivirus.

**Figure S3:**
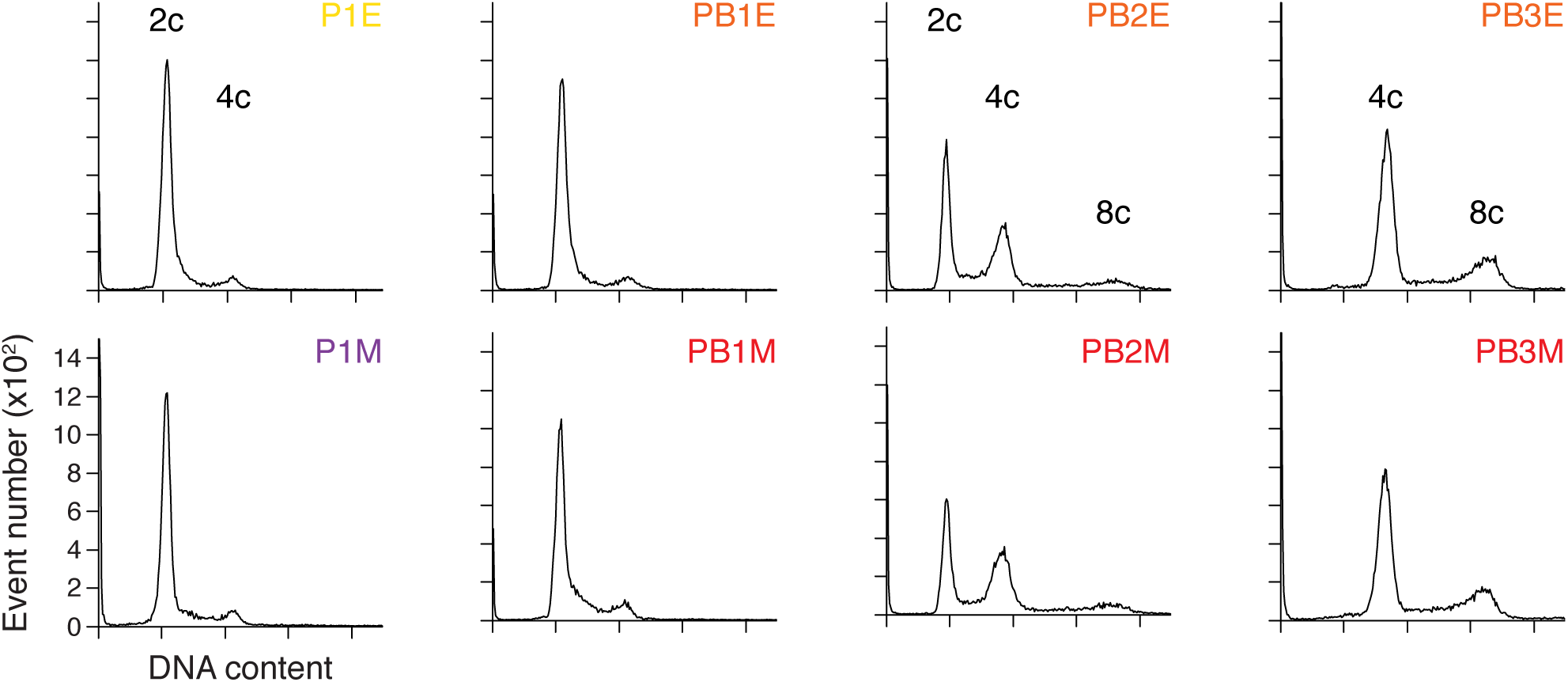
Genome Content of PB2 and PB3 Clones Suggests Aneuploidy. Flow cytometric analysis of genome content in control (empty vector) and MYC overexpressing cells of the same genotype. 2c, 4c and 8c correspond to a diploid, tetraploid and octoploid genome. P=*TP53*-mutant; B=*BRCA1*-mutant; E=empty vector lentivirus; M=*MYC*-overexpressing lentivirus.

**Figure S4:**
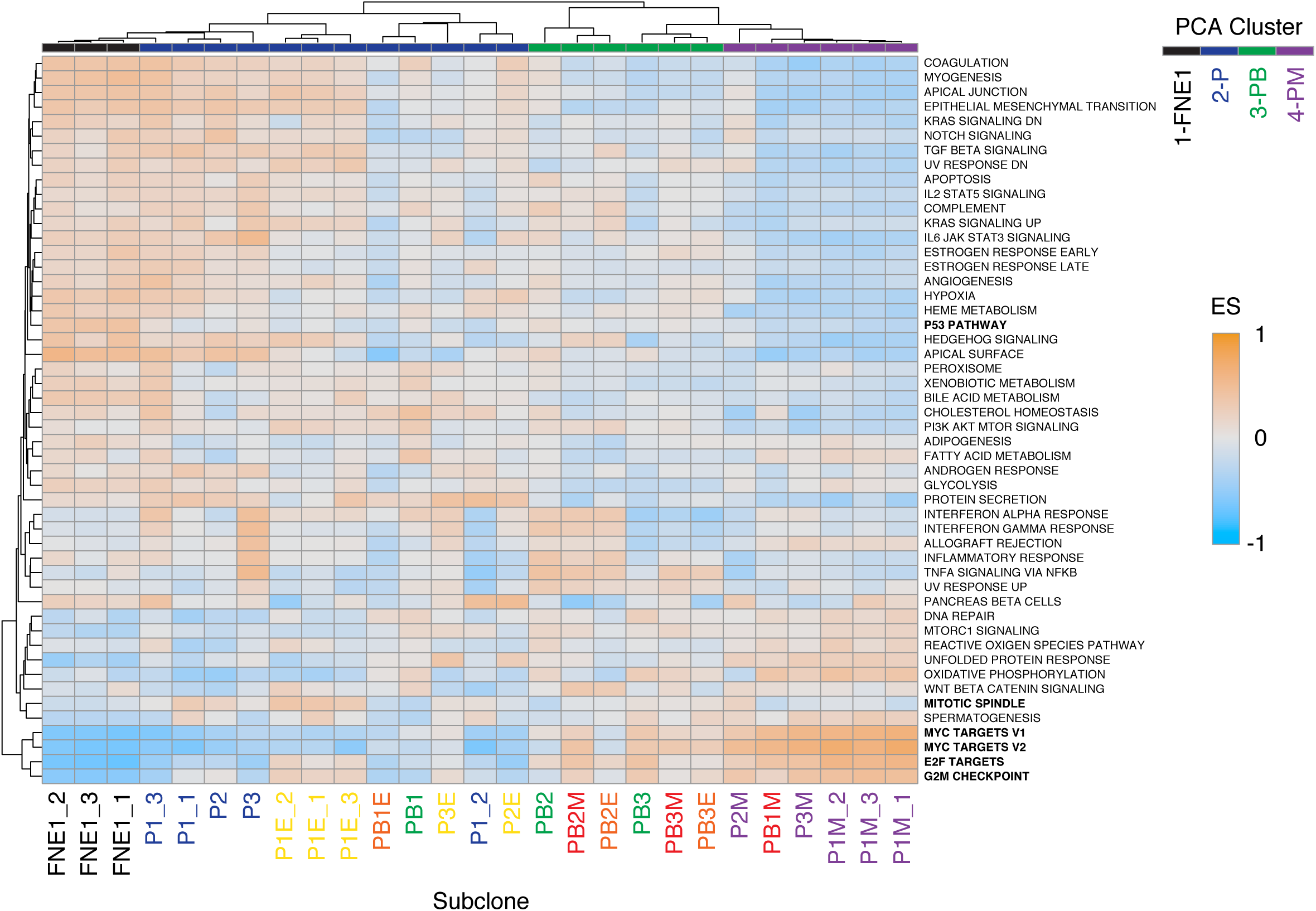
Gene Set Variation Analysis Separates Parental and Mutant Samples. Unsupervised hierarchical clustering of 27 cell lines based on enrichment scores calculated for Hallmark gene sets by gene set variation analysis (GSVA) from RNAseq. The top row indicates the PCA cluster of the respective sample, see Fig. 7A. Orange and blue shading indicate positive and negative enrichment scores, respectively.

**Figure S5:**
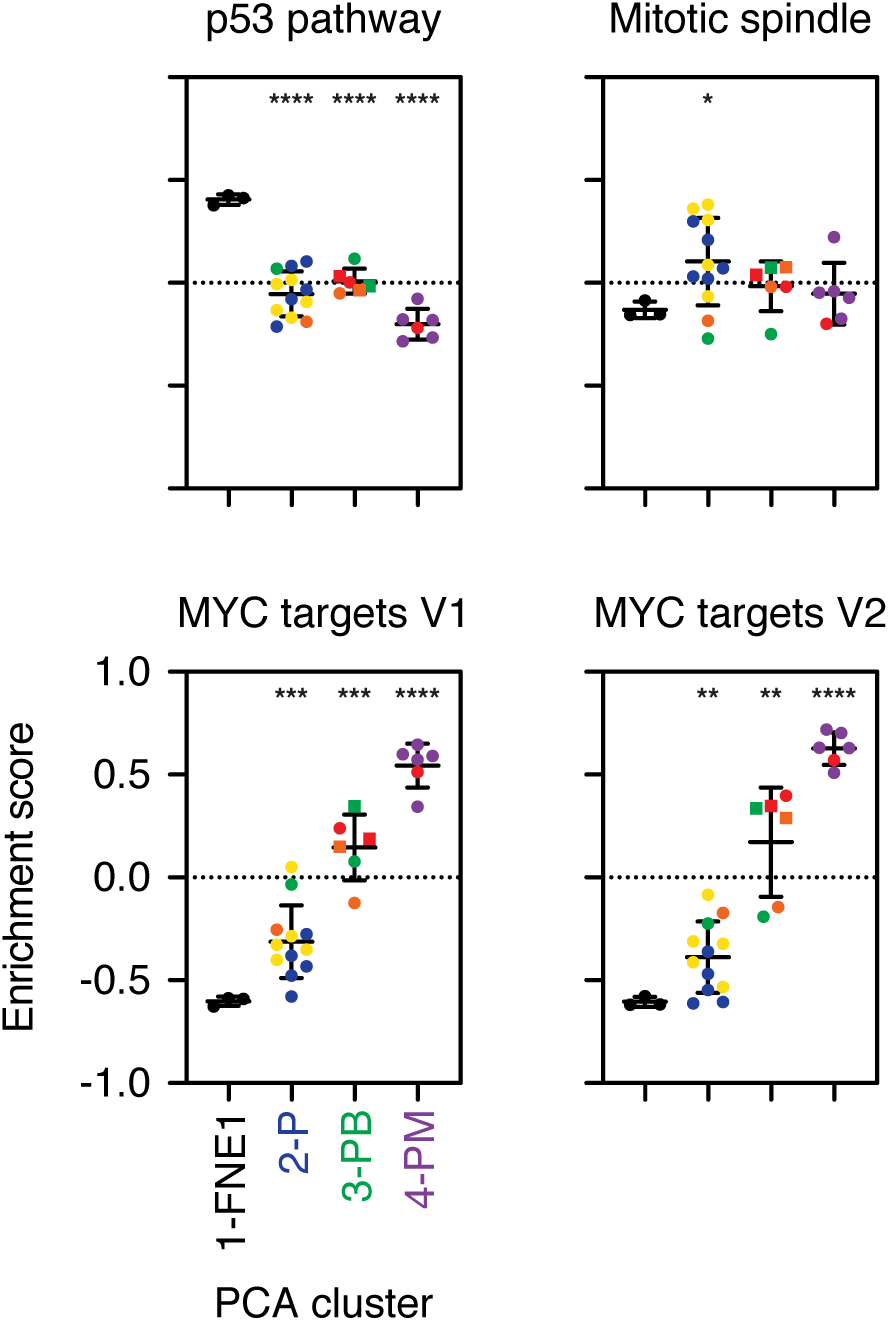
Gene Set Variation Analysis Corroborates Genotypic Transcriptomic Features. Results from four representative Hallmark gene sets from Fig. 6B are shown. Samples were grouped based on PCA cluster allocation and the colour of individual data points corresponds to sample genotype as in Fig. 6A. For cluster 1 (FNE1): n=3 samples; cluster 2 (P): n=12; and clusters 3 and 4 (PB and PM): n=6. Note PB1 and PB1E/M samples are included in clusters 2 and 4, respectively, rather than 3 (see text). Samples from the PB3 lineage are depicted as squares. Horizontal bar and error bars indicate mean and standard deviation, respectively. Asterisks depict adj. p-value between indicated groups compared with cluster 1 (FNE1) by Brown-Forsythe and Welsh ANOVA where * adj. p-value ≤ 0.05, ** adj. p-value ≤ 0.005, *** adj. p-value ≤ 0.0005, **** adj. p-value < 0.0001. See Table S5.

**Figure S6:**
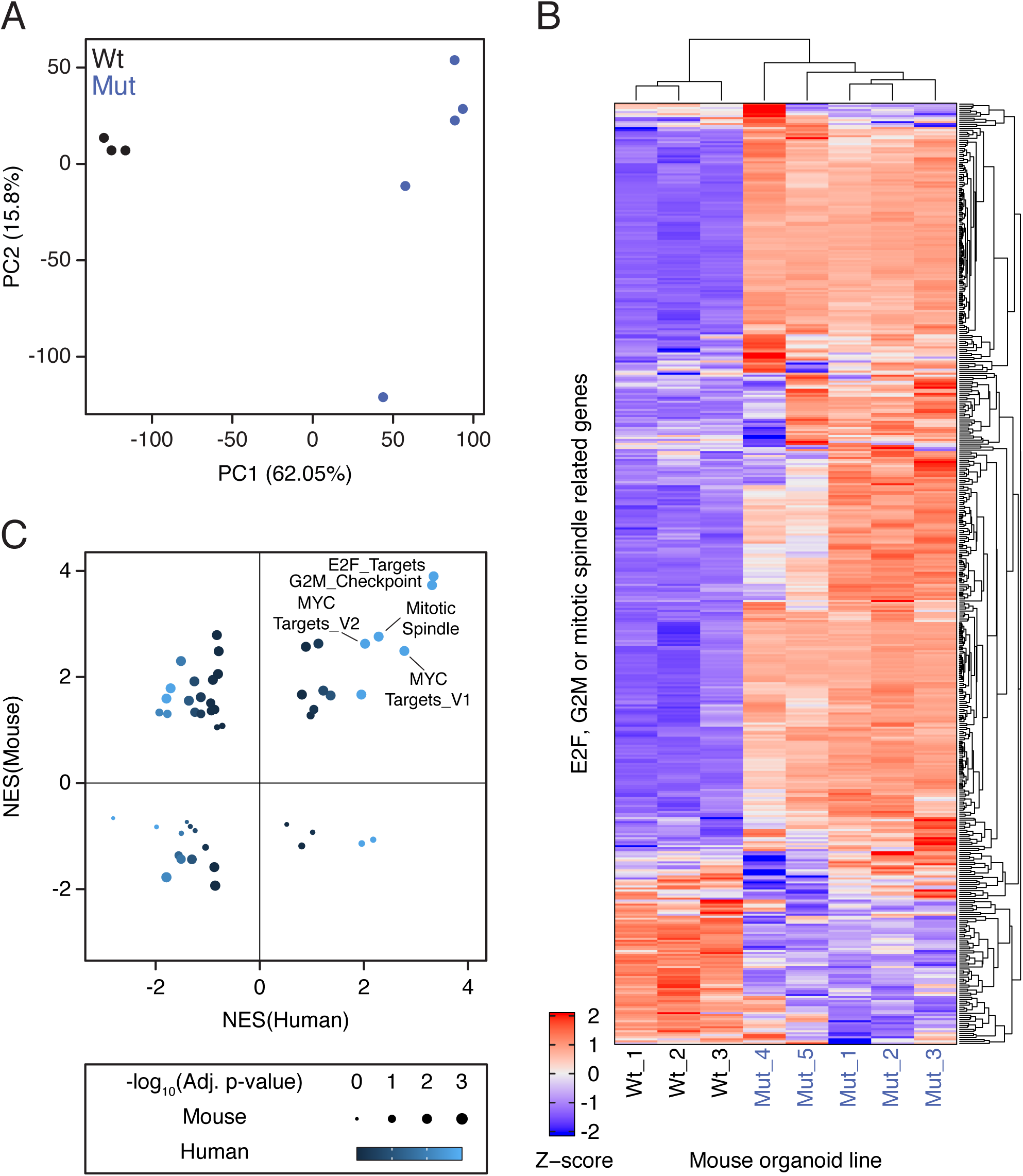
Differential Expression of Cell Cycle Regulators in *TP53-*mutant Mouse Fallopian Tube Organoids Correlates with that of Human *TP53*-mutant Fallopian Tube-derived Subclones. **A** Principal component analysis (PCA) of publicly available RNA sequencing data from eight murine wildtype (Wt) and *Trp53*-mutant (Mut) organoids (Zhang et al., 2019). Percent variance of principle components 1 (PC1) and 2 (PC2) are indicated in parenthesis along axes. See also Table S6. **B** Unsupervised hierarchical clustering based on the expression of 468 cell cycle regulators in the eight available mouse organoid samples. See also Table S7. **C** Correlation of positively and negatively enriched gene sets when *TP53* is mutated in our human FNE1 model and the *Trp53*-mutant mouse organoids versus corresponding control cells. The size and the colour of the bubbles indicate significance in the mouse and human contrasts with wildtype, respectively. NES=normalized enrichment score.

**Table S1**

Summary of reagents and critical commercial kits, experimental models and software.

**Table S2**

Summary of oligonucleotides used in this study. Blue font indicates gRNA sequence.

**Table S3**

Filtered, quantile normalized, batch corrected, Log_2_ transformed RNA sequencing reads for cell line samples used as basis for all human RNA sequencing analyses downstream used to generate Fig. 7.

**Table S4**

Mean enrichment scores for Hallmark gene sets calculated by gene set variation analysis of parental FNE1, P, PB and PM samples used to generate Fig. 7B.

**Table S5**

Enrichment scores calculated in gene set variation analysis (GSVA) of all samples used to generate data in Table S4, Fig. 7B,C, S4, S5.

**Table S6**

Filtered, quantile normalized, batch corrected, Log_2_ transformed RNA sequencing reads for organoid samples used as basis for all mouse RNA sequencing analyses downstream used to generate Fig. S6.

**Table S7**

Z-scores calculated sample-wise for mouse organoid samples used to generate Fig. S6B.

## Notes

### Competing Interest Statement

The authors have declared no competing interest.

